# Chromatin dynamics associated with sexual differentiation in a UV sex determination system

**DOI:** 10.1101/2020.10.29.359190

**Authors:** Josselin Gueno, Simon Bourdareau, Guillaume Cossard, Olivier Godfroy, Agnieszka Lipinska, Leila Tirichine, J. Mark Cock, Susana M. Coelho

## Abstract

In many eukaryotes, such as dioicous mosses and many algae, sex is determined by UV sex chromosomes and is expressed during the haploid phase of the life cycle. In these species, the male and female developmental programs are initiated by the presence of the U- or V-specific regions of the sex chromosomes but, as in XY and ZW systems, phenotypic differentiation is largely driven by autosomal sex-biased gene expression. The mechanisms underlying sex-biased transcription in XY, ZW or UV sexual systems currently remain elusive. Here, we set out to understand the extent and nature of epigenomic changes associated with sexual differentiation in the brown alga *Ectocarpus*, which has a well described UV system. Five histone modifications, H3K4me3, H3K27Ac, H3K9Ac, H3K36me3, H4K20me3, were quantified in near-isogenic male and female lines, leading to the identification of 13 different chromatin states across the *Ectocarpus* genome that showed different patterns of enrichment at transcribed, silent, housekeeping or narrowly-expressed genes. Chromatin states were strongly correlated with levels of gene expression indicating a relationship between the assayed marks and gene transcription. The relative proportion of each chromatin state across the genome remained stable in males and females, but a subset of genes exhibited different chromatin states in the two sexes. In particular, males and females displayed distinct patterns of histone modifications at sex-biased genes, indicating that chromatin state transitions occur preferentially at genes involved in sex-specific pathways. Finally, our results reveal a unique chromatin landscape of the U and V sex chromosomes compared to autosomes. Taken together, our observations reveal a role for histone modifications in sex determination and sexual differentiation in a UV sexual system, and suggest that the mechanisms of epigenetic regulation of genes on the UV sex chromosomes may differ from those operating on autosomal genes.

## Introduction

In species that reproduce sexually, sex is often determined by a pair of sex chromosomes: X and Y chromosomes in male-heterogametic species, Z and W in female-heterogametic species or U and V in haploid sexual systems (Bachtrog et al., 2014). Sex chromosomes originate from pairs of autosomes, but become differentiated after the sex-specific chromosome (Y, W or both the V and U) stops recombining (Bachtrog et al., 2014; Charlesworth, 2017; Umen and Coelho, 2019). Males and females have distinct sex chromosome sets but the extensive phenotypic differences between males and females (sexual dimorphism) are largely caused by differences in autosomal gene expression, so-called sex-biased gene expression. The nature and extent of sex-biased gene expression has been investigated in recent years across a broad range of taxa using genome-wide transcriptional profiling. These studies have revealed that sex-biased gene expression is common in many species, although its extent may vary greatly among tissues or developmental stages (reviewed in Grath and Parsch, 2016).

Although many reports have described patterns and evolution of sex-biased genes across several taxa, the molecular mechanisms underlying the regulation of sex-biased expression of hundreds, or even thousands, of genes during sexual differentiation remain poorly understood. One possible mechanism to regulate gene expression is through epigenetic modifications. Epigenetic modifications are defined as reversible changes that affect the genomic structure and regulate gene expression without affecting the DNA sequence itself (Allis and Jenuwein, 2016). Epigenetic modifications may occur through mechanisms such as DNA methylation and histone post-translational modifications (PTMs). DNA methylation regulates transcription in diverse eukaryotes (reviewed in Jones, 2012), and may contribute to transcriptional differences between sexes (Nugent et al., 2015), playing for instance an important role in differentiating female morphs (workers and queens) in the honeybee (Elango et al., 2009). In the liverwort *Marchantia*, male and female gametes have different levels of DNA methylation and this is correlated with differences in the expression of genes involved in DNA methylation (Schmid et al., 2018). Histone PTMs are another important component of transcriptional regulation, and can impact gene expression by altering chromatin structure or recruiting histone modifiers. Combination of histone PTMs (so-called chromatin states) are associated with functionally distinct regions of the genome such as heterochromatic regions and regions of either active transcription or repression (Kouzarides, 2007). The role of chromatin states in regulating gene expression patterns during development in animals is well established (Lindeman et al., 2010; Srivastava et al., 2010). However, very few studies have carried out chromatin profiling during sexual differentiation to link profiles with sex-biased expression patterns. The only available study, to our knowledge, described genome-wide maps of histone PTMs coupled with gene expression data to decipher the relationship between the chromatin states and sex-biased gene expression in *Drosophila miranda* (Brown and Bachtrog, 2014). In this study, the genome-wide distribution of both active and repressive chromatin states differed between males and females but sex-specific chromatin states appeared not to explain sex-biased expression of genes.

In organisms with XY or ZW sex determination systems, sex chromosomes often exhibit unique patterns of gene expression and unusual patterns of epigenetic marks compared with autosomes (e.g. Brown and Bachtrog, 2014; Schmid et al., 2018). For instance, in *Drosophila* males, where the Y is transcriptionally repressed and the X is hyper-transcribed (Baker et al., 1994), both of these transcriptional modifications are correlated with changes in the chromatin configuration (Gelbart and Kuroda, 2009; Girton and Johansen, 2008; Lemos et al., 2010; Straub and Becker, 2007). Sex chromosomes are derived from autosomes, but they are governed by unique evolutionary and functional pressures (Bachtrog, 2006). The sex-limited chromosome (Y or W) degenerates, i.e., loses most of its ancestral gene content, accumulates repetitive DNA and evolves a heterochromatic appearance (Bachtrog, 2013; Charlesworth and Charlesworth, 2000) whereas the homologous chromosome (X or Z) acquires mechanisms to compensate and evolves hyper-transcription (dosage compensation) (Lucchesi et al., 2005; Picard et al., 2018; Vicoso and Charlesworth, 2009). In *Drosophila* the euchromatin/heterochromatin ratio is different in the two sexes mainly due to the presence of the repeat-rich Y chromosome in males (Brown and Bachtrog, 2014; Yasuhara and Wakimoto, 2008). Similarly, the Z-specific region in schistosomes has a unique chromatin landscape, dominated by gene-activation-associated histone PTMs, that is associated with dosage compensation (Picard et al., 2019).

At present, no information is available concerning the regulation of gene expression by chromatin remodelling in organisms with UV sexual systems, such as mosses and algae (Coelho et al., 2018), although recent work has analysed the patterns of histone post translational modifications during the haploid-diploid life cycle of the brown alga *Ectocarpus* (Bourdareau et al., 2020). In UV sexual systems, sex is expressed during the haploid phase of the life cycle. Inheritance of a U or a V sex chromosome after meiosis determines whether the multicellular adult individual will be female or male, respectively (Bachtrog et al., 2014; Coelho et al., 2019). UV systems differ markedly from XY and ZW systems (Bull, 1978; Coelho et al., 2019; Umen and Coelho, 2019). For example, the two sexes are not homozygotic and heterozygotic so mechanisms such as chromosome-scale dosage compensation or meiotic sex chromosome inactivation are not expected. Moreover, whereas Y or W sex chromosomes often undergo genetic degeneration resulting in them being markedly different to their partner X or Z chromosome in terms of size, repeat content and gene density, U and V chromosomes do not tend to exhibit this type of asymmetry because each chromosome functions independently in a haploid context and therefore experiences similar evolutionary pressures (Ahmed et al., 2014).

The expression pattern of the genes located on U and V sex chromosomes has been shown to differ from that of the autosomal gene set (Coelho et al., 2019). For example, in the brown alga *Ectocarpus*, most sex-linked genes are upregulated during the haploid, gametophyte phase of the life cycle (Ahmed et al., 2014; Lipinska et al., 2017). The pseudo-autosomal regions (PARs) of the sex chromosomes are enriched in both life cycle-related genes (sporophyte-biased genes) and female-biased genes, compared to the autosomes (Lipinska et al., 2015). Moreover, PAR genes display unusual structural features compared with autosomal genes in terms of their GC content, repeat content and intron sizes (Avia et al., 2018; Luthringer et al., 2015).

Here, we investigated the sex-related chromatin landscape of *Ectocarpus*, a model brown alga with a UV sexual system. Comparison of the profiles of five histone PTMs with transcriptomic data showed that chromatin states were predictive of transcript abundance. The chromatin state of genes that exhibited sex-biased expression was markedly different in males and females indicating that histone modifications may play an important role in mediating sexual differentiation. Moreover, an important subset of the PAR genes presented sex-specific chromatin patterns. The U and V sex chromosomes were found to have very different chromatin landscapes to autosomes, despite the absence of a requirement for chromosome-scale dosage compensation in *Ectocarpus* and the fact that the U and V chromosomes do not exhibit strong signs of genetic degeneration.

## Results

### Identification of chromatin states in males and females of *Ectocarpus* sp

Near-isogenic male and female gametophyte lines (Table S1, Figure S1) were used to generate sex-specific chromatin immunoprecipitation and sequencing (ChIP-seq) profiles for five different histone PTMs: H3K4me3, H3K9ac, H3K27ac, H3K36me3 and H4K20me3 (Table S2). H3K4me3 is a near-universal chromatin modification that has been found at the transcription start sites (TSS) of expressed genes in a range of eukaryotes, and is associated with gene transcription (Barski et al., 2007; He et al., 2010; Howe et al., 2017). H3K9ac is a chromatin mark that is often associated with ongoing transcription in both animals and land plants (Brusslan et al., 2015; Heintzman et al., 2007). H3K27ac is an important mark that can distinguish between active and poised enhancer elements in animals (Creyghton et al., 2010). H3K36me3 is a gene body mark associated with active gene transcription in animals and plants (Roudier et al., 2011; Shilatifard, 2006). H4K20me3 is a repressive, constitutive heterochromatin mark but also silences repetitive DNA and transposons. H4K20me3 is generally associated with heterochromatin but its presence at gene bodies has been inversely correlated with gene expression in animals (Nelson et al., 2016; Schotta et al., 2004).

Given the large phylogenetic distances separating the brown algae from the animal and land plant lineages and the independent evolution of multicellularity in each of these three lineages (Cock et al., 2010), it is possible that the five histone PTMs analysed here are not associated with the same functions in brown algae as they are in animals and land plants. However, a previous analysis of histone PTMs in *Ectocarpus*, which included the marks tested here (Bourdareau et al., 2020), afforded evidence for similar roles. Peaks of H3K9ac, H3K27ac and H3K4me3 were detected within 500 bp of transcription start sites (TSSs). H3K36me3 and H4K20me3 were depleted from TSSs and transcription end sites (TESs), being associated with gene bodies and H4K20me3 was also present in intergenic regions. Together, the five histone PTMs used in our study are therefore expected to provide a broad overview of the *Ectocarpus* sp. chromatin landscape in male and female algae.

Thirteen chromatin states (i.e., different combinatorial patterns of histone PTMs) were defined in the *Ectocarpus* genome based on analysis of the genome-wide distribution patterns of the five histone PTMs using MACS2 (Zhang et al., 2008) and SICER (Xu et al., 2014) (Figure 1A). States S9-S13 consisted of combinations of histone marks that are usually associated with active transcription (presence of H3K36me3, H3K27ac, H3K9ac, H3K4me3) (Bourdareau, 2018). States S2-S8 all included H4K20me3, in most cases in addition to one or more of the above gene activation-associated marks. State S1 corresponded to a ‘background’ state, i.e., domains that were not enriched for any of the histone PTMs assayed. An example of histone PTM profiles for a 20 kbp region of the *Ectocarpus* genome is shown in Figure 1B.

**Figure 1.**
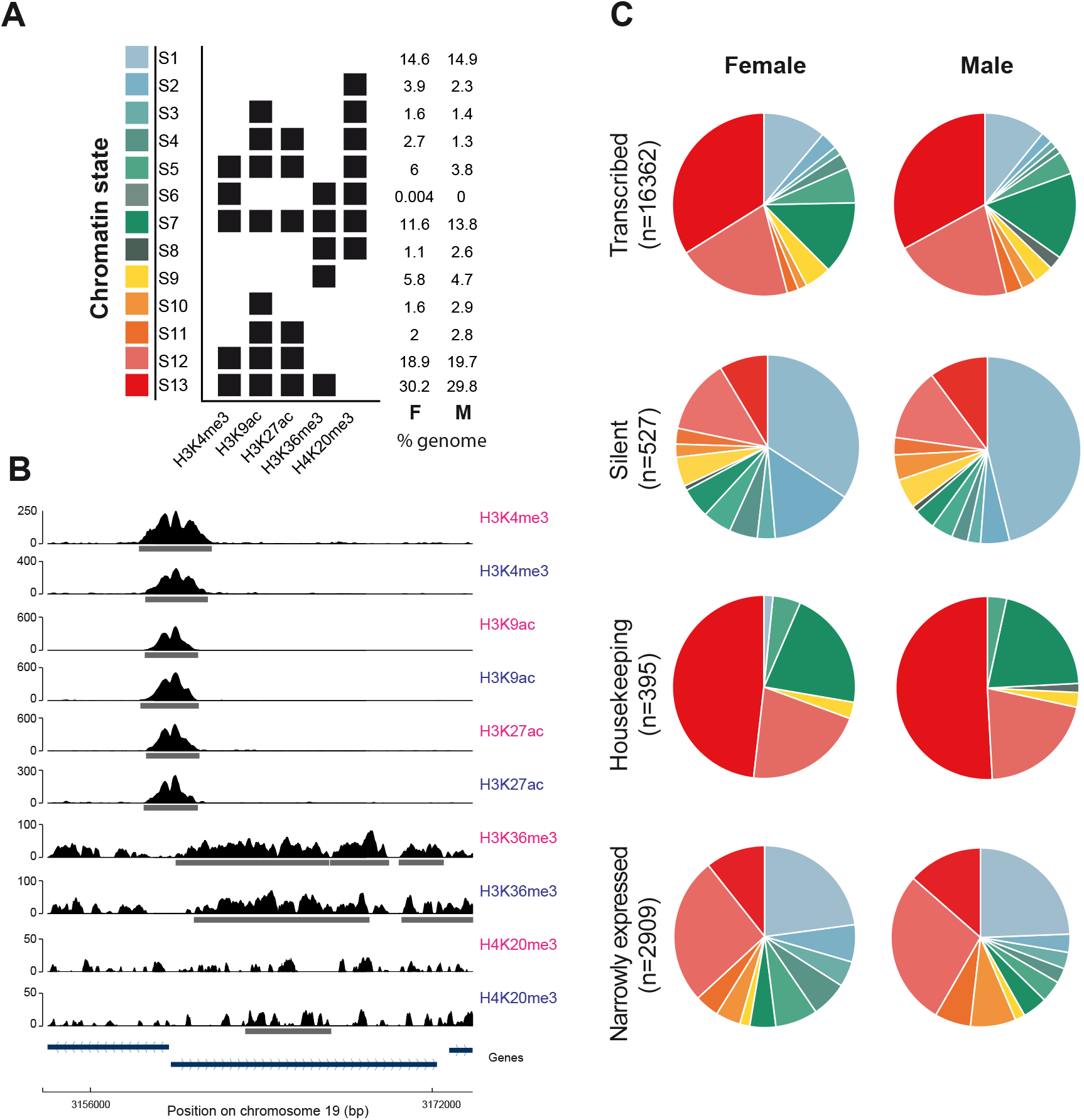
The chromatin landscape of male and female *Ectocarpus* sp. A) Summary of the 13 chromatin states detected in *Ectocarpus* sp. Percentages of the total gene set associated with each chromatin state in males (M) and females (F) are shown to the right. B) Representative region of the chromosome 19 showing profiles of mapped ChIP-seq reads for the five histone PTMs in males and females. Grey bars represent the peaks detected by MACS2 (H3K4me3, H3K9ac and H3K27ac) or SICER (H3K36me3 and H4K20me3). Blue bars represent genes. Pink text, females; blue text, males. C) Proportions of transcribed (TPM≥5th percentile), silent (TPM<5th percentile), housekeeping (tau<0.25) and narrowly expressed genes (tau>0.75) associated with each chromatin state in males and females.

### Chromatin states of different categories of *Ectocarpus* genes

To elucidate the relationship between the observed chromatin states and the expression patterns of *Ectocarpus* sp. genes, RNA-seq data was generated using the same biological samples as were used for the ChIP-seq analysis (see methods) and these data, together with previously published datasets (Lipinska et al., 2015, 2017, 2013), were used to define four categories of genes based on their expression patterns: transcribed genes (TPM≥5th percentile), silent genes (TPM<5th percentile), housekeeping genes (i.e. broadly expressed genes defined as having values of less than 0.25 for the tissue specificity index tau; see methods) and narrowly expressed genes (tau>0.75; see methods). The housekeeping and narrowly expressed genes (NEGs) were subsets of the transcribed gene set.

The most common chromatin state for the transcribed genes (32.9% and 33.9% in males and females respectively) was S13, which corresponds to co-localisation of all four of the histone PTMs that are generally associated with gene activation (H3K36me3, H3K27ac, H3K9ac, H3K4me3; Figure 1C, Table S3). For the ‘silent’ category of genes, S1 (no detectable histone PTM peak) was the most common state (45.4% and 34.0% in males and females, respectively; Figure 1C, Table S3). Housekeeping (broadly expressed) genes and NEGs have been shown to have distinct patterns of chromatin PTMs in *Drosophila* (Brown and Bachtrog, 2014; Filion et al., 2010). The majority (50.9% and 48.2% in males and females, respectively) of the housekeeping genes in *Ectocarpus* were associated with state S13 (all four marks associated with activation) whereas NEGs exhibited no clearly preferred state, the most common state being S12 (H3K4me3, H3K9ac, H3K27ac; 28.2% and 26.2% in males and females, respectively; Figure 1C, Table S3). States that included H4K20me3 were more common at NEGs than at housekeeping genes. Conversely, states associated with H3K36me3 (S6-S9 and S13) were characteristic of housekeeping genes (i.e., 75.9% and 72.3% of the housekeeping genes in males and females respectively had H3K36me3), and this mark was distinctly less prevalent on NEGs (19.8% and 17.1% in males and females respectively; Figure 1C, Table S3, Table S4). Finally, the background state S1 (none of the tested marks associated) was markedly more frequent at NEGs than at housekeeping genes. Together, these data support the association of the tested marks with active or repressed chromatin states in *Ectocarpus*.

When the relative proportions of the chromatin states were compared between males and females for each of the four gene categories (transcribed, silent, housekeeping and narrowly expressed), broadly similar patterns were observed in the two sexes, but some small differences were also noticeable (Figure 1C, Table S3). For example, less than 1% of the transcribed genes corresponded to state S8 (i.e., combination of H3K36me3 and H4K20me3) in females, compared to 2.4% in males (Table S3). Also, state S6 (combination of H3K4me3, H3K36me3 and H4K20me3) was exclusively present in a small subset of genes in females. Taken together, these results indicate that overall, the relative proportions of the different chromatin states across the genome remain relatively stable in males versus females.

### Identification of histone PTMs associated with gene activation and gene repression

To further investigate the relationship between the observed chromatin states and gene expression, transcript abundances in both males and females were plotted for the sets of genes corresponding to each chromatin state. A clear trend towards increasingly higher levels of transcript abundance was correlated with the gradual acquisition of the histone PTMs H3K9ac, H3K27ac, H3K4me3 and H3K36me3 (in the following order: H3K9ac followed by H3K9ac/H3K27ac, then by H3K9ac/H3K27ac/H3K4me3 and finally by H3K9ac/H3K27ac/H3K4me3/H3K36me3; Figure 2A; Table S5, S6). These observations support the proposed association of these four histone PTMs with gene activation (Bourdareau et al., 2020). These results also suggest that there may be a hierarchy in terms of the deposition of these histone PTMs, with addition of later marks being dependent on the presence of earlier ones in the order H3K9ac, H3K27ac, H3K4me3 and H3K36me3.

**Figure 2.**
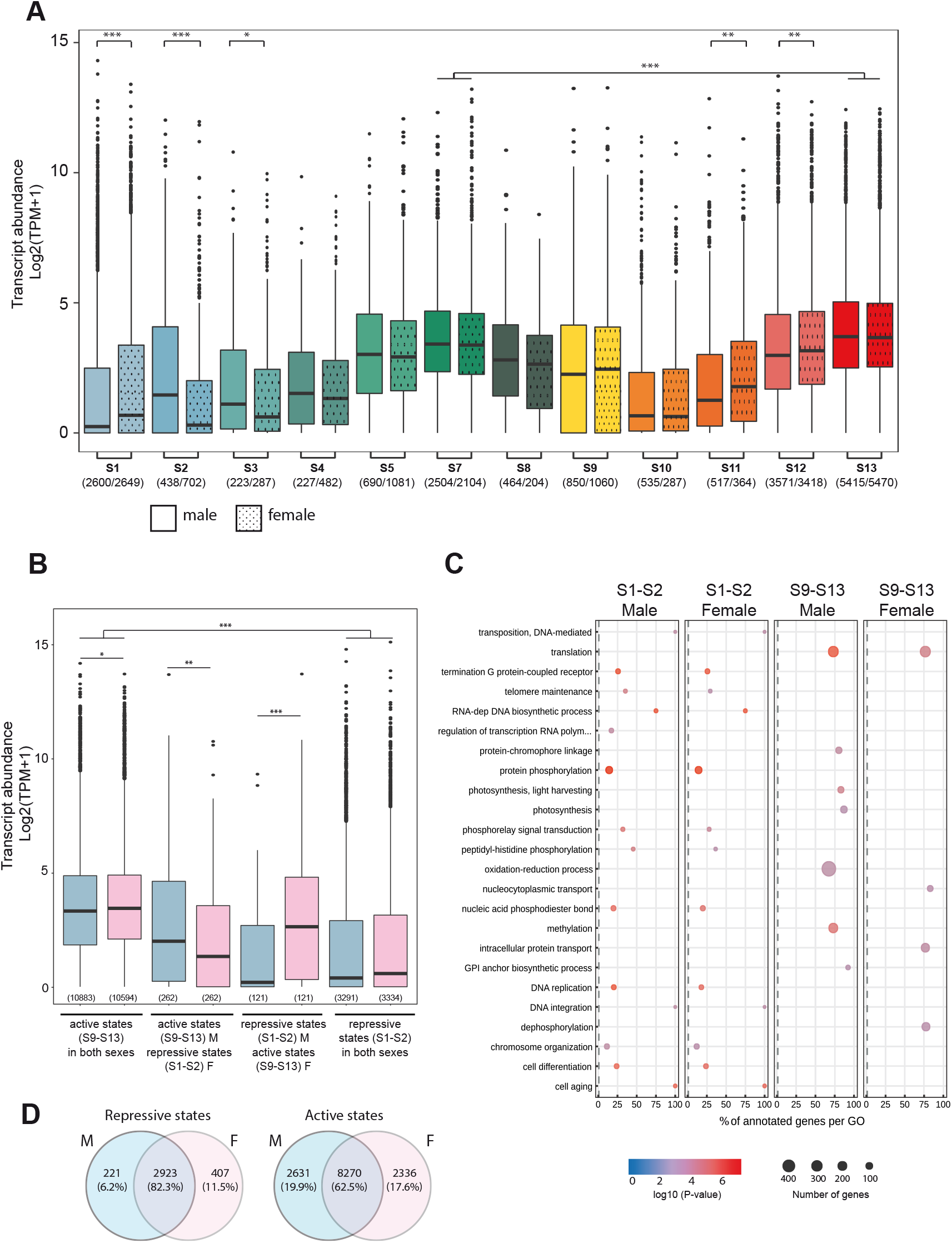
Gene expression and chromatin states. A) Transcript abundances for genes associated with different chromatin states in males and females. The colour code is the same as that used in Figure 1A. The numbers of genes associated with each state are indicated in brackets (males/females). Asterisks above plots indicate significant differences in gene expression (pair-wise Wilcoxon test, *p-value<0.05, **p-value<0.01, ***p-value<0.001). The full set of statistical tests is presented in Table S6. B) Transcript abundances for genes exhibiting either activation-associated (S9 to S13) or repression-associated (S1 or S2) chromatin states in females (pink) and males (blue). Numbers in brackets indicate the number of genes in each class. Asterisks indicate significant differences (*p-value>0.05, **p-value<0.01, ***p-value<0.001). C) GO term enrichment for genes marked with activation-associated (S9-S13) or repression-associated (S1-S2) chromatin states in males and females. D) Venn diagrams representing the proportion of genes marked with activation-associated (S9-S13) or repression-associated (S1-S2) chromatin states in males and females.

In pairwise comparisons, sets of genes corresponding to chromatin states that included H4K20me3 consistently exhibited lower transcript abundance than sets of genes with equivalent chromatin states without H4K20me3 (e.g. transcript abundance was significantly lower for S7 than for S13; Wilcox test, p-value= 4.463E-18 Figure 2A; Table S5, S6). These results are consistent with H4K20me3 playing a role in the repression of gene expression in *Ectocarpus*. Note however that because H4K20me3 is frequently associated with transposons (Bourdareau et al., 2020), the observed association with transcriptional repression could also be indirect, via the silencing of intronic transposon sequences.

Finally, the background state S1 corresponds to domains that are not associated with any of the assayed histone PTMs, and *Ectocarpus* genes associated with state S1 exhibited very low transcript abundance (Figure 2A, Table S5, S6).

Analysis of the RNA-seq data also indicated some differences between the sexes. For example, on average, genes in chromatin state S1, S11 and S12 had significantly higher expression levels in females compared with males (pairwise Wilcoxon, p-value=2.4E-7; p-value=0.02 and p-value=0.001, respectively; Figure 2A). Conversely, on average, genes in chromatin state S2 and S3 had lower expression levels in females than in males (pairwise Wilcoxon, p-value=6.3E-8, p-value=3.4E-8; Figure 2A, Table S5, S6).

To further examine the link between chromatin states and transcript abundances in males and females, we classified states S1 and S2 (absence of any of the tested marks or presence of only H4K20me3) as ‘repressive’ chromatin states, while states S9-S13 were classified as ‘active’ chromatin states (presence of at least one canonical activation-associated mark H3K9ac, H3K27ac, H3K4me3 and/or H3K36me3). Note that we did not include genes with states S3-S7 in this analysis because they exhibited a combination of repression-associated (H4K20me3) and activation-associated marks and because they were expressed at intermediate levels (Figure 2A). As expected, genes marked with states S9-S13 were expressed at higher levels in both sexes than those that were associated with states S1 and S2 (Figure 2B; pair-wise Wilcox test, p-value<2E-16). Interestingly, levels of gene expression in males and females were also significantly different for genes marked with states S9-S13 in one sex but with states S1 and S2 in the other (Figure 2B; pair-wise Wilcox test, p-value<2E-16). Therefore, sex-specific differences in the chromatin states of genes were associated with sex-specific expression patterns.

A GO-term enrichment analysis of genes in either activation-associated states (S9-S13) or repression-associated states (S1-S2) showed that the set of genes associated with states S9-S13 was enriched in functions such as translation, oxidation-reduction, methylation and dephosphorylation, whereas the set of genes in S1-S2 states was enriched in functions such as phosphorylation and DNA replication (Figure 2C). GO term enrichment was more stable between sexes for repression-associated chromatin states, whereas sex-specific GO term enrichment was observed for genes in the activation-associated chromatin states S9-S13 (Figure 2C, Table S7) but note that a large proportion of the genes S1-S2 exhibited conservation of the repression-associated state in both males and females (82.3% of genes), whereas conservation was less marked for genes in states S9-S13 (62.5%; Figure 2D).

### Chromatin states and gene expression of *Ectocarpus* sex-biased genes

To investigate the role of histone PTMs in sexual differentiation, we examined the chromatin states associated with genes that showed sex-biased expression patterns. A comparison of gene expression patterns in the two near-isogenic male and female lines (Figure S1), based on RNA-seq data generated using the same biological samples as were used for the ChIP-seq analysis, identified a total of 268 genes that exhibited sex-biased expression (padj<0.05, fold change>2, TPM>1; Table S5).

Presence of the activation-associated chromatin marks H3K36me3, H3K9me3, H3K27ac and H3K4me3 (states S9-S13) was associated with higher transcript abundance for sex-biased genes in both males and females but the difference was only statistically significant for males (Wilcoxon test p-values of 0.012 for males and 0.188 for females; Figure S2, S3). Sex-biased genes therefore display a similar association between the presence of activation-associated marks and increased gene expression levels as observed with the genome-wide gene set (Figure 2A).

### Chromatin states of sex-biased genes in males and females

To analyse modifications of chromatin PTMs associated with differential expression of sex-biased genes, transitions between chromatin states in males and females were evaluated on a gene-by-gene basis. This analysis showed that 54.8% of male-biased genes (MBGs) and 47.2% of female-biased genes (FBGs) had different chromatin states in males and females (Table S5), underlining the dynamic landscape of histone PTMs on sex-biased genes in males and females.

Overall, the proportions of the different chromatin states, specifically for MBGs, were significantly different compared with NEGs suggesting that their chromatin landscape is not related to their narrow expression (Chi-square test, p-value = 4.937E-15 and p-value = 0.01608 in FBGs vs NEGs in females and males and p-value = 5.627E-4 and p-value = 3.333E-6 for MBGs vs NEGs in females and males respectively; Figure 1C and Figure 3A).

**Figure 3.**
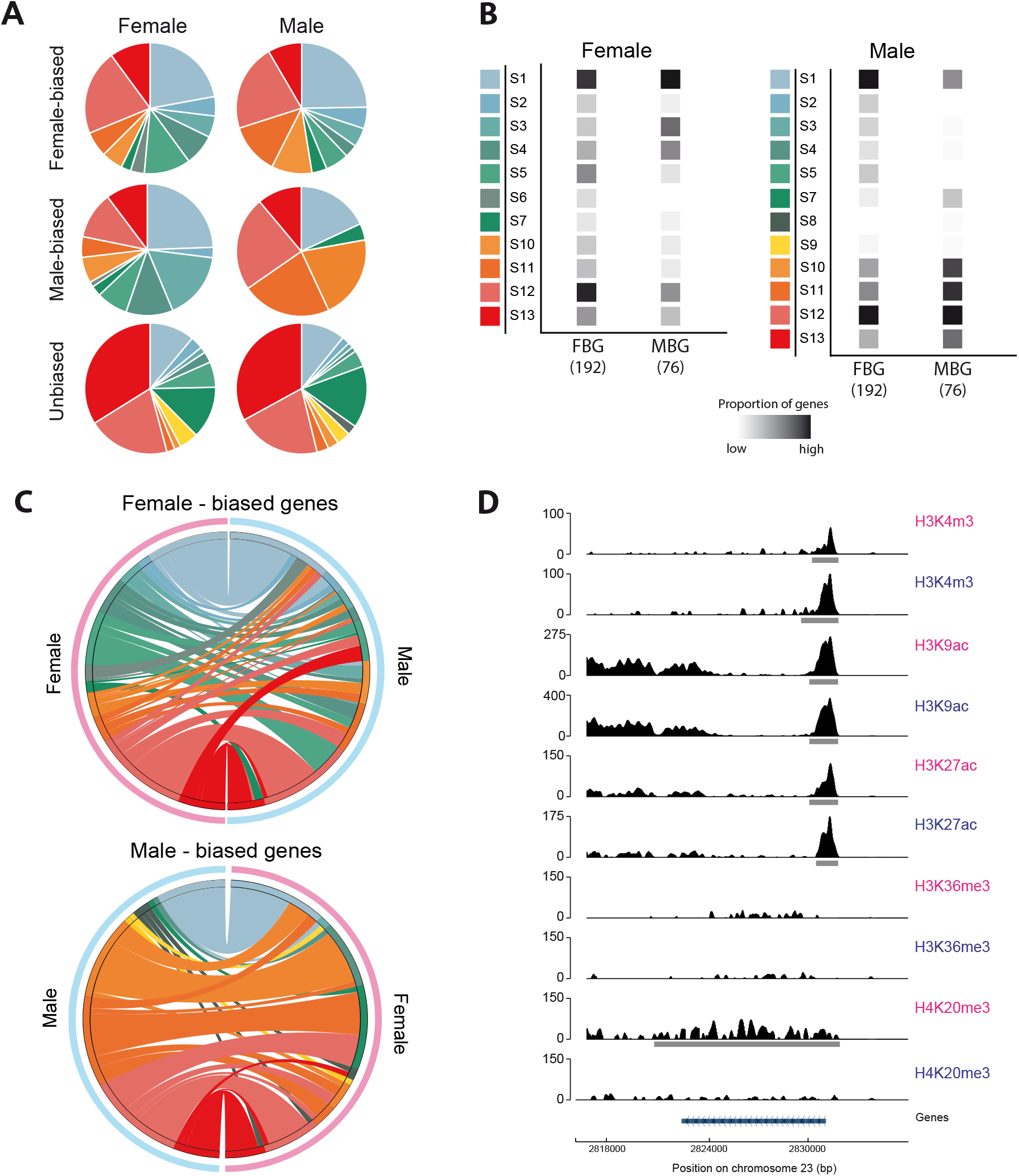
Histone PTM patterns at sex-biased genes in *Ectocarpus* sp. males and females. A) Proportions of the 13 chromatin states for female-biased, male-biased and unbiased genes in females (left) and males (right). B) Proportions of genes associated with each of the 13 chromatin states for female-biased and male-biased genes in females (left) and males (right). The intensity of the grey squares is proportional to the number of genes corresponding to each state. Coloured squares represent the different chromatin states (see Figure 1A). The total numbers of genes analysed for each condition is given in brackets, and the number of genes in each chromatin state are provided in Table S8. FBG: female-biased genes; MBG: male-biased genes. C) Circos plots comparing chromatin states associated with female-biased (above) and male-biased (below) genes in females (pink) and males (blue). The colour code for the chromatin states is the same as that used in Figure 1A. Each link corresponds to the transition from a state in the sex on the left to a state in the sex on the right of the circos plot. D) Representative chromatin profiles for a male-biased gene on chromosome 23 (blue bar). The histone PTMs indicated in blue and pink correspond to those of the male and the female, respectively. The horizontal grey bars under each track correspond to peaks called by either MACS2 (H3K4me3, H3K9ac and H3K27ac) or SICER (H3K36me3 and H4K20me3).

For the set of male-biased genes there was a marked difference between the relative proportions of the different chromatin states in males compared to females: in males, chromatin states that included the repression-associated mark H4K20me3 were rare whereas states that included activation-associated marks (H3K9ac, H3K27ac, H3K4me3 and/or H3K36me3, but not H4K20me3) were common (Figure 3A, 3B; Table S3). The proportion represented by chromatin states that contained H4K20me3 (S2 to S8) decreased from 42.1% in females to 3.9% in males whilst the proportion represented by states S9 to S13 (activation-associated states) increased from 34% in females to 73.7% in males (Figure 3A, 3B; Table S3). Almost half (43.4%) of the male-biased genes exhibited a transition from a state that included H3K9ac, H3K27ac, H3K4me3 and/or H3K36me3, but not H4K20me3 (S9-S13) in males to a state that either included H4K20me3 (S2 to S8) or to state S1 (none of the histone PTMs detected) in females (Figure 3C; Table S9). The chromatin state transitions of male-biased genes were consistent with the correlation between the presence and absence of activation-associated and repression-associated histone PTMs and differences in the abundances of the transcripts of sex-biased genes between sexes observed for the complete set of all *Ectocarpus* genes (Figure 2A).

Unexpectedly however, female-biased genes exhibited a different pattern of chromatin state transitions when males and females were compared. States that included activation-associated marks (H3K9ac, H3K27ac, H3K4me3 and/or H3K36me3, i.e. states S3-S13) were slightly more frequent in females (76.3%) compared with males (69.2%), but female-biased genes were often associated with H3K20me3 (states S2-S8) in females (37.8%; Figure 3A, B; Table S3). Only 12% of the female-biased genes were associated with chromatin marks S9-S13 in females and underwent a transition to a state that included H3K20me3 or to a background state in males (S1-S8) (Figure 3C; Table S9).

In conclusion, chromatin state transitions between sexes were concomitant with changes in expression levels of sex-biased genes between males and females. For male-biased genes, the patterns of these state transitions were consistent with the tendencies observed for the complete set of all *Ectocarpus* genes, and therefore with the associations between specific histone PTMs and either gene activation or gene repression reported for animals and land plants, as described above. Female-biased genes, however, did not conform to this pattern.

### The chromatin landscape of the *Ectocarpus* sex chromosomes

In organisms with diploid sexual systems (XY or ZW), sex chromosomes exhibit different patterns of histone PTMs to autosomes (Brown and Bachtrog, 2014; Picard et al., 2019). In *Drosophila* males for example, the X chromosome is transcribed at a higher level in males than in females, due to dosage compensation of the hemizygous X, and exhibits an enrichment in active chromatin marks (Brown and Bachtrog, 2014). In contrast, in female mammals, the inactivated X chromosome is characterized by DNA methylation and widespread presence of repressive chromatin marks (Brockdorff and Turner, 2015; Lucchesi et al., 2005). In addition, Z-chromosome-localised female-specific hyperacetylation of histone H4 (H4K16Ac) has been described for the chicken (Bisoni et al., 2005) and epigenetic analysis underlined the specialized chromatin landscape of the Z-specific region of *S. mansooni*, which is more permissive than that of the autosomal regions in both male and female *S. mansooni* (Picard et al., 2019).

A similar marked difference between sex chromosomes and autosomes was observed in *Ectocarpus* sp. (Figure 4A, Table S5, S10; Figure S4, S5). The relative proportions of each of the 13 chromatin states showed some variance between autosomes but the set of genes on the sex chromosomes exhibited strikingly different patterns to those of the autosomes (Figure 4A). There was a significant dearth of genes marked with the activation-associated states S12 and S13 on the sex chromosomes compared to the autosomes (permutation tests U versus autosomes, p-value_S12_=0.047 and p-value_S13_=0.039; permutations tests V versus autosomes, p-value_S12_=0.046 and p-value_S13_=0.037; Table S11). Furthermore, in males, the sex chromosome was significantly enriched in states that included the histone PTM H4K20me3 compared with autosomes, specifically state S2 (p-value = 0.025), S4 (p-value = 0.021), S5 (p-value *=* 0.008) and S8 (p-value *=* 0.028); Figure 4A-C, Table S11).

**Figure 4.**
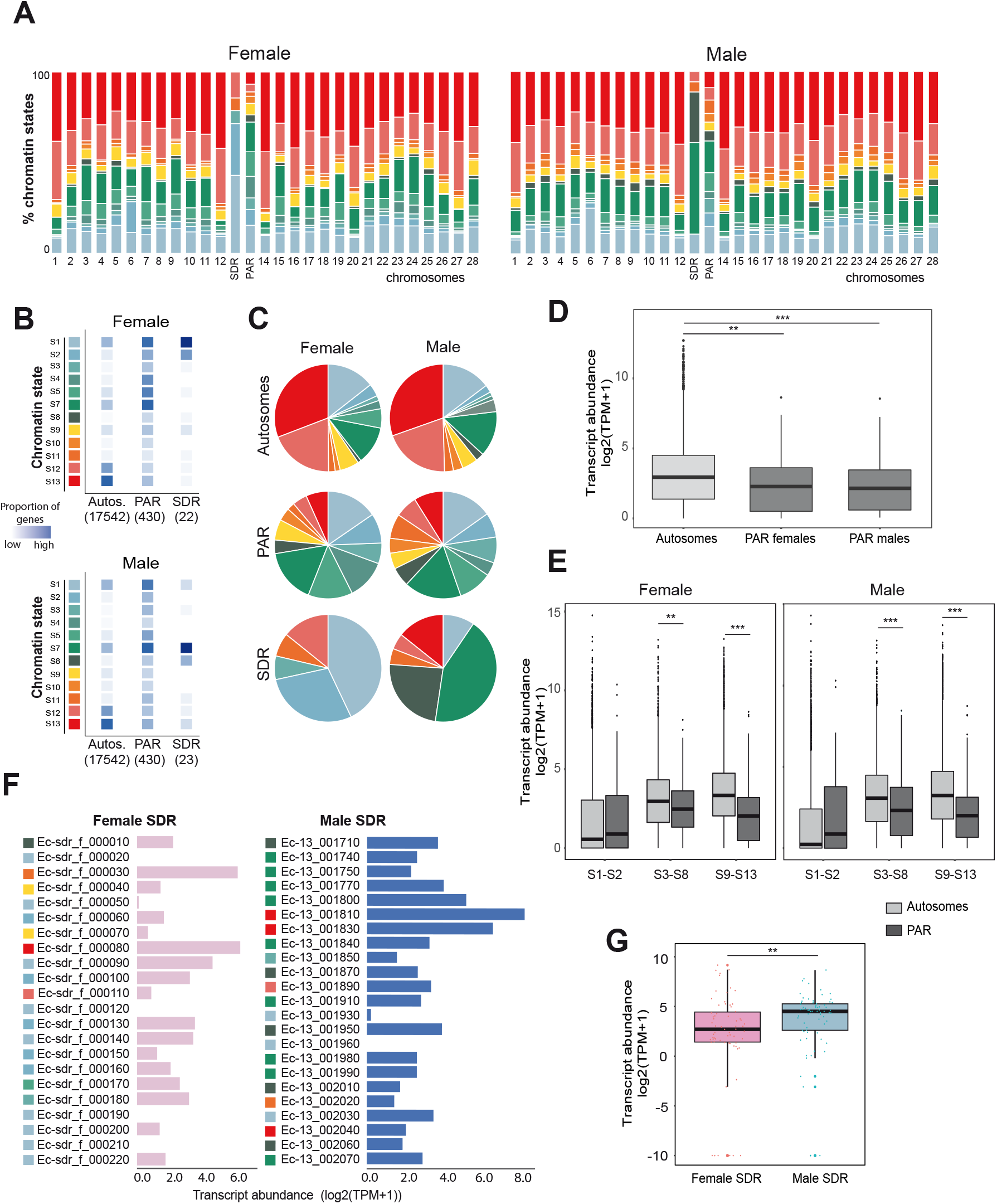
Chromatin landscape of the U and V sex chromosomes compared with the autosomes. A) Chromatin state distribution for each autosome and for the SDR and PAR regions of the sex chromosome in females (left panel) and in males (right panel). B) Proportions of genes associated with each of the 13 chromatin states for all autosomes and for the PAR and SDR regions of the sex chromosome in females (top panel) and in males (bottom panel). The intensity of the blue colour is proportional to the number of genes in each state. The total numbers of genes in each genomic region are represented in brackets. The colour code for the chromatin states is the same as that used in Figure 1A. Autos., autosomes. C) Proportions of chromatin states associated with autosomal, PAR and SDR genes in males and females. The colour code for the chromatin states is the same as in Figure 1A. D) Transcript abundances, measured as log2(TPM+1), for autosomal and for PAR genes in males and females. E) Transcript abundances for autosomal and PAR genes in different chromatin states: repression-associated states S1-S2, states that include canonical activation marks and H4K20me3 (S3-S8) and activation-associated states S9-S13. Significant differences were assessed using pairwise Wilcoxon rank sum test (**p-value < 0.01, ***p-value < 0.0001). F) Transcript abundances, measured as log2(TPM+1), for individual genes located in the female (pink) and male (blue) sex determining regions (SDRs). Coloured squares represent chromatin states corresponding to the colour code indicated in Figure 1A (see also Table S13). G) Transcript abundances of genes located within the sex-specific regions (SDRs) of the U and V sec chromosomes. Asterisks above the plots indicate significant differences (pair-wise Wilcoxon test, **p-value < 0.01).

The significantly distinct chromatin patterns between sex chromosome and autosomes were equally manifest when only the pseudoautosomal region (PAR) was taken into account (Chi-square test p-value <2.2E-16; Figure 4A-C). For example, 67% and 76% of the PAR genes in males and females, respectively, were associated with chromatin states S1-S8 compared with 40.1% and 41.5% in males and females, respectively, for autosomal genes (Table S5, Table S12). Although the proportions of chromatin states for the PAR in males and females were not statistically different (chi-square test with continuity correction, p-value=0.251) there were considerably more genes with chromatin state S4 in the PAR in females (12%) than in the PAR in males (4%) (Proportion test, p-value = 6.214e-06) (Table S12). Remarkably, almost half (45%) of the genes located in the PAR were found to be associated with different chromatin states in males and females (Table S5), indicating that a substantial proportion of the PAR genes display sex-dependent chromatin state transitions. Note that only 11 of the 412 PAR genes were classed as sex-biased genes (Table S5), so the sex-related changes in chromatin states of the PAR genes do not appear to be linked with sex-biased PAR gene expression.

Analysis of the sex-determining regions of the chromosomes showed that the majority of the genes within the female SDR (i.e., U-specific genes) were in state S1 (i.e., carried none of the assayed marks) whereas the V-specific genes were mostly in state S7 (displayed all of the assayed marks) or state S8 (H3K36me3 and H4K20me3), with some genes in state S13 (all marks except H4K20me3) (Figure 4B-C; Table S13). However, note that, due to the low number of SDR genes, it was not possible to rule out that the difference between chromatin state patterns of the male and female SDRs was due to chance (100 000 permutations tests on Pearson’s X^2^ statistics).

Previous work has shown that the *Ectocarpus* PAR region is enriched in transposons compared with autosomes (Ahmed et al., 2014; Luthringer et al., 2015). Considering that in *Ectocarpus* H4K20me3 co-localizes with transposon sequences (Bourdareau et al., 2020) we asked if the presence of transposons in PAR genes could explain the observed chromatin state distribution patterns. More PAR genes contained a transposon sequence compared to autosomal genes (80% versus 36%, respectively) but there was not a correlated increase in the proportion of PAR genes marked with H4K20me3 (28-29% for the PAR versus 25-27% for autosomes) (Table S14). Moreover, permutation tests using subsets of autosomal genes in which 80% of the genes were selected to contain transposons (i.e., a similar proportion of genes with transposons to that observed for the PAR) indicated that the unusual pattern of chromatin of states in the PAR was not due simply to the presence of additional genes with inserted transposons (Table S14).

Overall, transcript abundances of genes located in the PAR were significantly lower than those located in autosomes (Wilcoxon p-value=0.003 and p-value=0.0005 for female and male respectively; Table S4, Figure 4D). Potentially, this difference in expression level may also have explained the difference between the chromatin state patterns of the PAR and the autosomes. To test this hypothesis, we selected a subset of autosomal genes that had a similar pattern of transcript abundances to that of the PAR genes (Table S15). The distribution of chromatin states for this set of autosomal genes was different to that of the PAR genes (Figure S6, indicating that gene expression level was not the cause of the difference in chromatin state patterns between the PAR genes and the autosomes.

The lower transcript abundance for PAR genes was consistent with the higher proportion of genes in repressive-associated states (S1-S2) compared with autosomal genes (24% for the PAR compared with 13% for the autosomes), but note that even PAR genes in activation-associated states (S9-S13) exhibited significantly lower expression levels than autosomal genes in similar states (pairwise Wilcoxon test, p-value=7.1E-9, p-value=7.1E-9 for female and male respectively; Figure 4E).

### Chromatin states and expression levels of sex chromosome genes

Gene expression levels and deposition of chromatin marks were highly correlated for the complete set of *Ectocarpus* genes (see above, Figure 2A). For example, genes in state S13 (presence of all four activation-associated marks) had a significantly higher expression level compared with genes in state S7 (presence of all four activation-associated marks plus H4K20me3). In females, when the correlations between chromatin states and levels of gene expression were compared for the autosomes and for the PAR, three chromatin states (S7, S12 and S13) exhibited a significantly weaker correlation with expression for the latter compared with the former. In males, weaker correlation between chromatin state and expression level was also observed for the PAR compared to the autosomes but only for states S7 and S13 (Table S16, Figure S7). In other words, depending on the location (PAR or autosomes) the correlation between chromatin state and gene expression level was not the same.

There was no significant correlation between levels of expression of either male or female SDR genes and the presence of particular chromatin marks (likelihood ratio tests, p-value = 0.460 and p-value = 0.304 for female and male SDR, respectively; Figure 4F), but the small sample size of SDR genes decreases the power of the statistical test. Note however that H3K36me3, a mark associated with transcript elongation (Huang and Zhu, 2018), was more often present at male SDR genes (in 18/23 genes) than at female SDR genes (1/22 genes)(Figure 4, Figure S4, Table S5) and we also noticed that abundances of transcripts for male SDR genes were significantly higher than for female SDR genes (Figure 4G; pairwise Wilcoxon test with Holm correction, p-value=0.0098).

Taken together, these observations suggest that the sex chromosome exhibits exceptional features in terms of its chromatin landscape. The unique features of the PAR are not explained by the preponderance of intragenic transposons nor by the fact that genes in this region have a lower mean level of gene expression. The relationship between chromatin state and gene expression level for the sex chromosomes is different to that observed for the autosomes.

## Discussion

### Epigenetic regulation in a haploid UV sexual system

Three types of genetic sex determination system exist in nature: XX/XY, ZZ/ZW systems and U/V systems (Bachtrog et al., 2014; Coelho et al., 2018). Studies have focused on understanding sex determination and sex biased gene expression but we know little about chromatin dynamics in males compared to females. The objective of this study was to provide an overview of the sex differences in the chromatin landscape in a haploid UV system, and to investigate the relationship between chromatin states and gene expression differences between sexes and genomic regions, with a particular emphasis on the U and V sex chromosomes.

We analysed the genome-wide distribution of five histone PTMs in males and females of an organism with haploid UV sex determination, resulting in the definition of 13 chromatin states corresponding to different combinations of the five histone PTMs. Chromatin states that included different combinations of H3K4me3, H3K9ac, H3K27ac and H3K36me3 were associated with actively transcribed genes, whereas chromatin states that included H4K20me3 were associated with a decrease in gene expression compared to equivalent states that lacked H4K20me3. States that included H3K36me3 were associated with broadly expressed genes, and this mark was less prevalent on genes with narrow expression patterns, a configuration that is compatible with the idea that H3K36me3 is deposited during transcription elongation (Barski et al., 2007). Note that the difference in H3K36me3 levels in NEGs versus housekeeping genes could be related to the lower power to detect H3K36me3 binding of tissue-specific genes expressed only in a subset of cells. It was interesting however that the difference between the housekeeping and NEG gene sets was considerably more marked for H3K36me3 than for the TSS-located PTM (Table S4), perhaps indicating a stronger link with gene transcription. A similar association of H3K36me3 with broadly expressed genes has been described for *Drosophila* (Brown and Bachtrog, 2014; Filion et al., 2010), indicating that this correlation has been conserved across distantly related lineages. Overall, the *Ectocarpus* chromatin patterns described here are consistent with H3K4me3, H3K9ac, H3K27ac and H3K36me3 having similar roles in brown algae, land plants and animals (Baroux et al., 2011; Bourdareau et al., 2020; Margueron and Reinberg, 2010; She and Baroux, 2015). The role of H4K20me3, in contrast, appears to be less conserved across eukaryotic supergroups, being associated with low transcriptional levels in both animals and brown algae but with euchromatin and transcriptional activation in land plants (de la Paz Sanchez and Gutierrez, 2009; Fischer et al., 2006).

Our analysis has also identified some novel features of the relationship between chromatin marks and gene expression in *Ectocarpus*. For example, we identified a positive correlation between the number of different activation-associated marks (TSS marks and H3K36me3) that were deposited at a gene and transcript abundance. In the absence of canonical repressive marks such as H3K27me3 in *Ectocarpus* (Bourdareau et al., 2020), it is possible that chromatin regulation of gene expression in *Ectocarpus* may be dominated by the synergistic action of activation marks (although it is important to bear in mind the possibility that the activation-associated marks may be deposited a consequence of transcription rather than mediating gene activation). Deposition of H4K20me3 was consistently associated with decreased transcript abundance in *Ectocarpus*, and in that respect this mark can be considered to be ‘repression-associated`. However, it is currently unclear if H4K20me3 action is direct or indirect through silencing of intronic transposons (Bourdareau et al., 2020).

### Relationship between H4K20me3 and gene expression

A complex relationship was observed between H4K20me3 and gene expression. There was clear evidence for a correlation between H4K20me3 and gene expression levels that was independent of the TSS-localised marks and H4K36me3 (Figure 2A). A previous study found that genes marked with H4K20me3 exhibited significantly weaker signals for TSS-localised PTMs (H3K4me2, H3K4me3, H3K9ac, H3K14ac and H3K27ac) (Bourdareau et al., 2020), suggesting a possible effect of H4K20me3 on gene expression via TSS marks. Taken together, these observations suggest that H4K20me3 may act on gene expression via two different pathways, one via an effect on TSS marks and the other by acting directly on gene expression, independently of the TSS marks. However, an alternative hypothesis would be that increased gene expression leads to a decrease in H4K20me3. In other words, activation of a gene might involve (in addition to other processes) suppression of heterochromatin-associated marks such as H4K20me3 leading to a tendency for H4K20me3 to be present at loci that are less marked with TSS-located PTMs.

### Chromatin dynamics of *Ectocarpus* sex-biased genes in males and females

Genome-wide, the proportions of genes associated with each chromatin state did not differ substantially when males were compared with females. However, when individual genes were compared, a considerable fraction was associated with different chromatin states in the two sexes, including genes that did not exhibit sex-biased expression patterns. It is possible that the differences correspond to chromatin state ‘noise’, in which case they would not be expected to be linked with sex-biased gene expression. However, the strong correlation between chromatin states and expression levels argues for a biological role for chromatin state changes. One hypothesis would be that genes display sex-specific chromatin configurations prior to the appearance of significant sex differences in gene expression and phenotypic differentiation. In other words, differences in chromatin state may anticipate sex-biased differences in gene expression at later stages, as has been reported for mammalian fetal germ cells (Lesch and Page, 2013). A more refined study using several stages of development of male and female gametophytes would be needed to gain further insights into this matter.

In males, most of the male-biased genes were marked with activation-associated chromatin states (S9-S13), whereas in females, male-biased genes were predominantly marked with repression-associated chromatin states (S1-S2). This observation is consistent with gene expression level modifications reported for sex-biased genes in males compared with females, where male-biased expression is due to a combination of both upregulation in males (i.e., activation of male-biased genes in males) and decreased expression in females (i.e., repression of male-biased genes in females) (Lipinska et al., 2015). However, more than half of the FBGs were marked with activation-associated chromatin states (S9-S13) in males, whereas in females, FBGs were predominantly marked with chromatin states that included H4K20me3. It appears therefore that female-biased genes do not follow the same trends that were observed genome-wide and for MBGs, where TSS marks were clearly associated with gene activation and H4H20me3 associated with lower transcript abundances.

### Unique chromatin organisation features in the U and V sex chromosomes

In organisms with UV sexual systems, the U and V sex-specific regions are both non-recombining, exhibit relatively similar structural features and appear to have been subjected to similar evolutionary pressures (Ahmed et al., 2014; Mignerot and Coelho, 2016). Despite these similarities, the genes in the male SDR exhibited a different pattern of chromatin states to the genes in the female SDR. In particular, H3K36me3, a mark that is often involved in dosage compensation and is usually enriched on X chromosomes (Bell et al., 2008), was detected on 18/23 (78%) of the male SDR genes but only in 6% (1/16) female SDR genes, but note that statistical analysis showed no significant differences between U and V SDRs due to the low number of genes in this region. Deposition of H3K36me3 is associated with increased transcript abundances in plants and animals (Roudier et al., 2011; Shilatifard, 2006), and we found that genes on the *Ectocarpus* male SDR exhibited significantly higher expression levels than female SDR genes (Figure 4G).

The *Ectocarpus* PAR has been shown to have unusual structural and gene expression features compared to the autosomes (Avia et al., 2018; Luthringer et al., 2015) and this study found unusual patterns of chromatin states in this genomic region. However, the analysis also showed that neither the levels of gene expression, which are lower, on average, for the PAR compared with autosomes, nor the greater prevalence of transposons and repeat sequences in PAR genes explained the unusual patterns of chromatin states. Moreover, sex-specific differences in chromatin states were prominent on the PAR of the U and V sex chromosomes, where almost half (47%) of the genes displayed different chromatin states between the two sexes. Our observations emphasise the unique features of the PAR of the *Ectocarpus* UV sex chromosomes, and suggest that the effect of chromatin states on transcript abundance may depend on the genomic locations of genes, and that the same chromatin states do not correspond to the same level of transcriptional change in genes located in autosomes and sex chromosomes. It is possible that the expression of genes on the U and V sex chromosomes is regulated by different epigenetic processes to those that regulate the expression of autosomal genes, perhaps involving histone PTMs that have not been assayed in this study. Further investigations employing additional histone PTMs marks will be needed to further understand the extraordinary features of these chromosomes.

## Methods

### Biological Material

The near-isogenic male (Ec457) and female (Ec460) *Ectocarpus* sp. lines (Table S1) were generated by crossing brother and sister gametophytes for either four or five generations, respectively (Ahmed et al., 2014). The resulting male and female strains, therefore, had essentially identical genetic backgrounds apart from the non-recombining SDR. Male and female gametophytes were cultured until near-maturity for 13 days as previously described (Coelho et al., 2012) at 13°C in autoclaved natural sea water supplemented with 300 μl/L Provasoli solution (PES), with a light:dark cycle of 12:12 h (20 μmol photons.m^−2^.s^−1^) using daylight-type fluorescent tubes. All manipulations were performed in a laminar flow hood under sterile conditions.

### Comparisons of male and female transcriptomes using RNA-seq

RNA for transcriptome analysis was extracted from the same duplicate male and female cultures as were used for the ChIP-seq analysis. For each sex, total RNA was extracted from a mix of 90 gametophytes each, using the Qiagen Mini kit (http://www.qiagen.com). RNA quality and quantity were assessed using an Agilent 2100 bioanalyzer, associated with Qubit2.0 Fluorometer using the Qubit RNA BR assay kit (Invitrogen, Life Technologies, Carlsbad, CA, USA), as described previously (Lipinska et al., 2015, 2017).

For each replicate sample, cDNA was synthesized using an oligo-dT primer. The cDNA was fragmented, cloned, and sequenced by Fasteris (CH-1228 Plan-les-Ouates, Switzerland) using an Illumina HiSeq 4000 set to generate 150-bp single-end reads. See Table S1 for RNA-seq accession numbers.

Data quality was assessed using FastQC (http://www.bioinformatics.babraham.ac.uk/projects/fastqc; accessed May 2019). Reads were trimmed and filtered using Cutadapt (Martin, 2011) with a quality threshold of 33 (quality-cutoff) and a minimal size of 30 bp.

Filtered reads were mapped to version v2 of the *Ectocarpus* sp. genome (Cormier et al., 2017a) using TopHat2 with the Bowtie2 aligner (Kim et al., 2013). More than 85% of the sequencing reads from each library could be mapped to the genome (Table S1).

The mapped sequencing data were then processed with featureCounts (Liao et al., 2014) to obtain counts for sequencing reads mapped to genes. Gene expression levels were represented as transcripts per million (TPMs). Genes with expression values below the fifth percentile of all TPM values calculated per sample were considered not to be expressed and were removed from the analysis. This resulted in a total of 18,462 genes that were considered to be expressed.

Differential expression analysis was performed with the DESeq2 package (Bioconductor) (Love et al., 2014). Genes were considered to be male-biased or female-biased if they exhibited at least a twofold difference (fold change; FC) in expression between sexes with a false discovery rate (FDR) < 0.05. A list of the sex-biased genes can be found in Table S5.

To calculate breadth of expression we employed the tissue-specificity index tau (Yanai et al., 2005) using published expression data from nine tissues or stages of the life cycle (female and male immature and mature gametophytes, mixed male and female gametophytes, partheno-sporophytes, upright partheno-sporophyte filaments, basal partheno-sporophyte filaments, diploid sporophytes) from *Ectocarpus* sp. (Cormier et al., 2017a; Lipinska et al., 2015, 2019, 2017; Luthringer et al., 2015). This allowed us to define broadly expressed (housekeeping) genes (with tau<0.25) and narrowly expressed genes (tau>0.75).

### Genome-wide detection of histone PTMs

Male versus female *Ectocarpus* sp. gametophyte ChIP-seq experiments were carried for H3K4me3, H3K9ac, H3K27ac, H3K36me3, H4K20me3, and three controls (an input control corresponding to sonicated DNA, histone H3 and immunoglobulin G monoclonal rabbit (IgG)) as in (Bourdareau, 2018). RNA-seq data (see above) was generated from the same samples, to ensure that the histone PTM and gene expression data were fully compatible. For ChIP-seq, 2.8 g (corresponding to 2800 individual gametophytes) of *Ectocarpus* tissue was fixed for five minutes in seawater containing 1% formaldehyde and the formaldehyde eliminated by rapid filtering followed by incubation in PBS containing 400 mM glycine. Nuclei were isolated by grinding in liquid nitrogen and in a Tenbroeck Potter in nuclei isolation buffer (0.1% triton X-100, 125 mM sorbitol, 20 mM potassium citrate, 30 mM MgCl2, 5 mM EDTA, 5 mM β-mercaptoethanol, 55 mM HEPES at pH 7.5 with complete ULTRA protease inhibitors), filtering through Miracloth and then washing the precipitated nuclei in nuclei isolation buffer with and then without triton X-100. Chromatin was fragmented by sonicating the purified nuclei in nuclei lysis buffer (10 mM EDTA, 1% SDS, 50 mM Tris-HCl at pH 8 with cOmplete ULTRA protease inhibitors) in a Covaris M220 Focused-ultrasonicator (duty 25%, peak power 75, cycles/burst 200, duration 900 seconds at 6°C). The chromatin was incubated with an anti-histone PTM antibody (anti-H4K20me3, anti-H3K4me3, and anti-H3K9ac, Cell Signal Technology; anti-H3K27ac, Millipore; anti-H3K36me3, Abcam) overnight at 4°C and the immunoprecipitation carried out using Dynabeads protein A and Dynabeads protein G. Following immunoprecipitation and washing, a reverse cross-linking step was carried out by incubating for at least six hours at 65°C in 200 mM NaCl and the samples were then digested with Proteinase K and RNAse A. Purified DNA was analysed on an Illumina HiSeq 4000 platform with a single-end sequencing primer over 50 cycles. At least 20 million reads were generated for each immunoprecipitation. The ChIP-seq dataset has been deposited in the NCBI Gene Expression Omnibus database under the accession numbers described in Table S2.

Quality control of the sequence data was carried out using FastQC (Andrews, 2016). Poor quality sequences were removed and the high quality sequences trimmed with Cutadapt (Hansen et al., 2016; Martin, 2011). Illumina reads were mapped onto the *Ectocarpus* genome (Cormier et al., 2017b) using Bowtie (Langmead et al., 2009). Duplicates were removed using samtools markdup in the Samtools package (v 1.9) (Li et al., 2009).

Quality control of ChIP-seq data sets followed the Encode ChIP-seq guidelines and practices (Landt et al., 2012)(Table S2). ChIP-seq analysis was carried out for two biological replicates for each PTM in both the male and female samples. Pearson correlation analysis of replicates was performed with multiBamSummary and then by plotCorrelation (v3.1.2 deepTools) (Ramirez et al., 2014). Replicate samples were strongly correlated (Pearson correlations >0.92, Figure S8).

To identify peaks and regions of chromatin mark enrichment, each data set, after combining data for biological replicates, was analysed separately for the male and female gametophyte. Peaks corresponding to regions enriched in H3K4me3, H3K9ac and H3K27ac were identified using the MACS2 (version 2.1.1) callpeak module (minimum FDR of 0.01) and refined with the MACS2 bdgpeakcall and bdgbroadcall modules (Zhang et al., 2008). H3K36me3 and H4K20me3 were analysed using SICER (v1.1) (minimum FDR of 0.01) (Xu et al., 2014; Zang et al., 2009) with a window size of 200 bp and a gap size of 400 bp. Note that peaks associated with sex-biased, PAR and SDR genes were manually inspected to validate reproducibility between replicates. The signal was normalized using the Signal Extraction Scaling (SES) method (Diaz et al., 2012).

Heatmaps, average tag graphs and coverage tracks were plotted using EaSeq (Lerdrup et al., 2016). Circos graphs were generated using Circos software (Krzywinski et al., 2009).

### Detection of chromatin states

To identify the patterns of histone PTM marks associated with each gene (i.e., chromatin states), we used bedtools intersect (intersectBed) from the Bedtools software (v2.26)(Quinlan and Hall, 2010). A total of 13 combinations of histone PTM marks (S1 to S13) were detected. Note that only chromatin states that were present in more than 1% of the genes were taken in consideration for the analysis.

### Coverage for each chromatin state

The coverage for each histone PTM per chromosome was calculated using bedtools coverage where the coverage of each PTM was normalized by the size of the chromosome. The pseudoautosomal regions (PAR) and the sex-specific, non-recombining regions (SDR) of the sex chromosome were analysed separately, as in (Brown and Bachtrog, 2014).

### Statistical analysis

To test for significant differences in the conservation of chromatin states between sex-biased and unbiased genes, we used mixed generalised linear models with a binomial distribution, modelling conserved vs non-conserved states as a function of bias. We then performed a likelihood ratio test with a null model to assess the significance of bias. Statistical analysis was performed in R 3.6.3. Permutation tests were performed to study the differences of proportions of chromatin states in PAR and SDR genes compared to autosomal genes. We randomly subsampled 100,000 times a number of chromatin states equal to the number of PAR genes, SDR genes or both, from autosomal genes in order to perform proportion tests. We compared observed and simulated Pearson’s chi-square statistics to assess whether the observed differences in chromatin state proportions between gene sets (autosomal, SDR, PAR, SDR+PAR) were statistically due to chance. A significant p-value indicates that the observed difference in proportion are not due to chance. In order to eliminate any possible effect of TE prevalence (which is different between PAR, SDR and autosomal genes) we also performed these tests using a randomized set of autosomal genes that displayed exact the same TE prevalence.

### GO-term analysis

Gene set enrichment analysis (GSEA) was carried out separately for each sex and each histone state, using Fisher’s exact Test implemented in the R package TopGO using the weight01 algorithm to account for GO topology (Alexa and Rahnenfuhrer, 2020) We investigated enrichment in terms of molecular function ontology and report significant GO-terms with p-value < 0.01.

## Supplemental Figures

**Figure S1.**
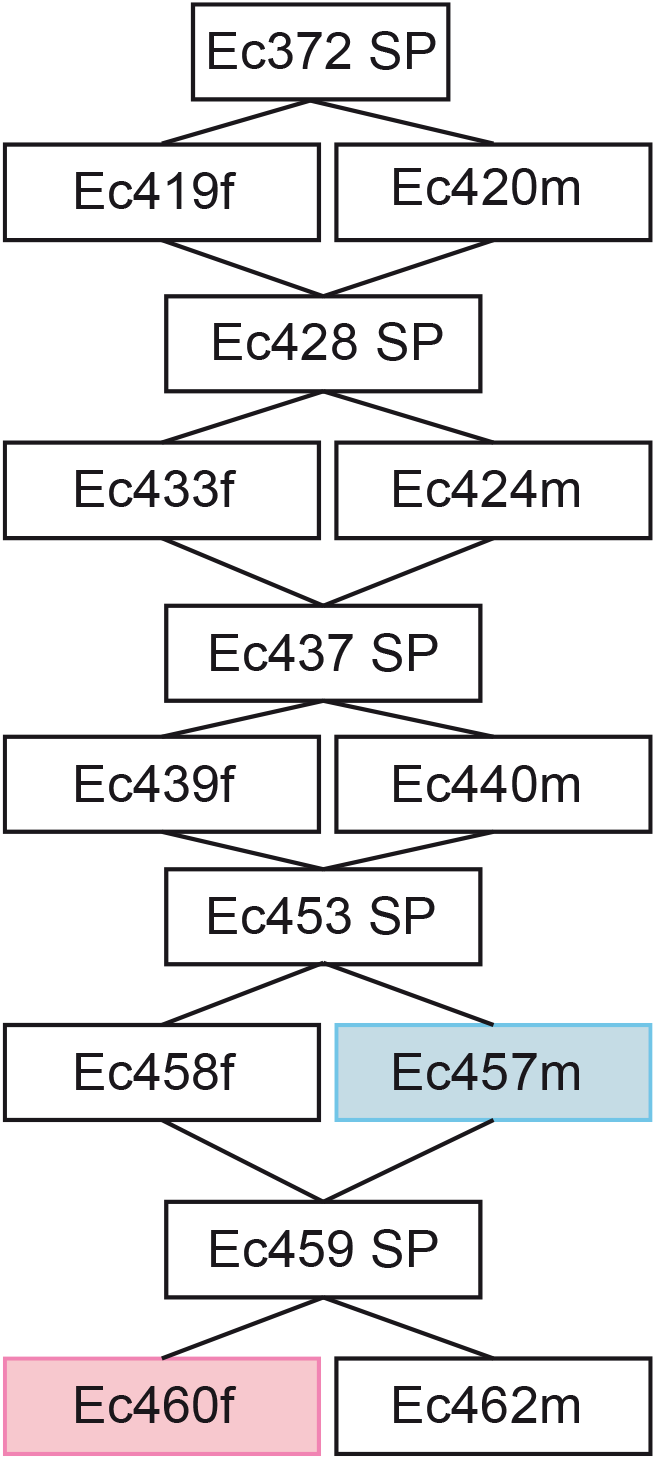
Pedigree of the male and female strains used in this study. SP, sporophyte; m, male gametophyte; f, female gametophyte.

**Figure S2.**
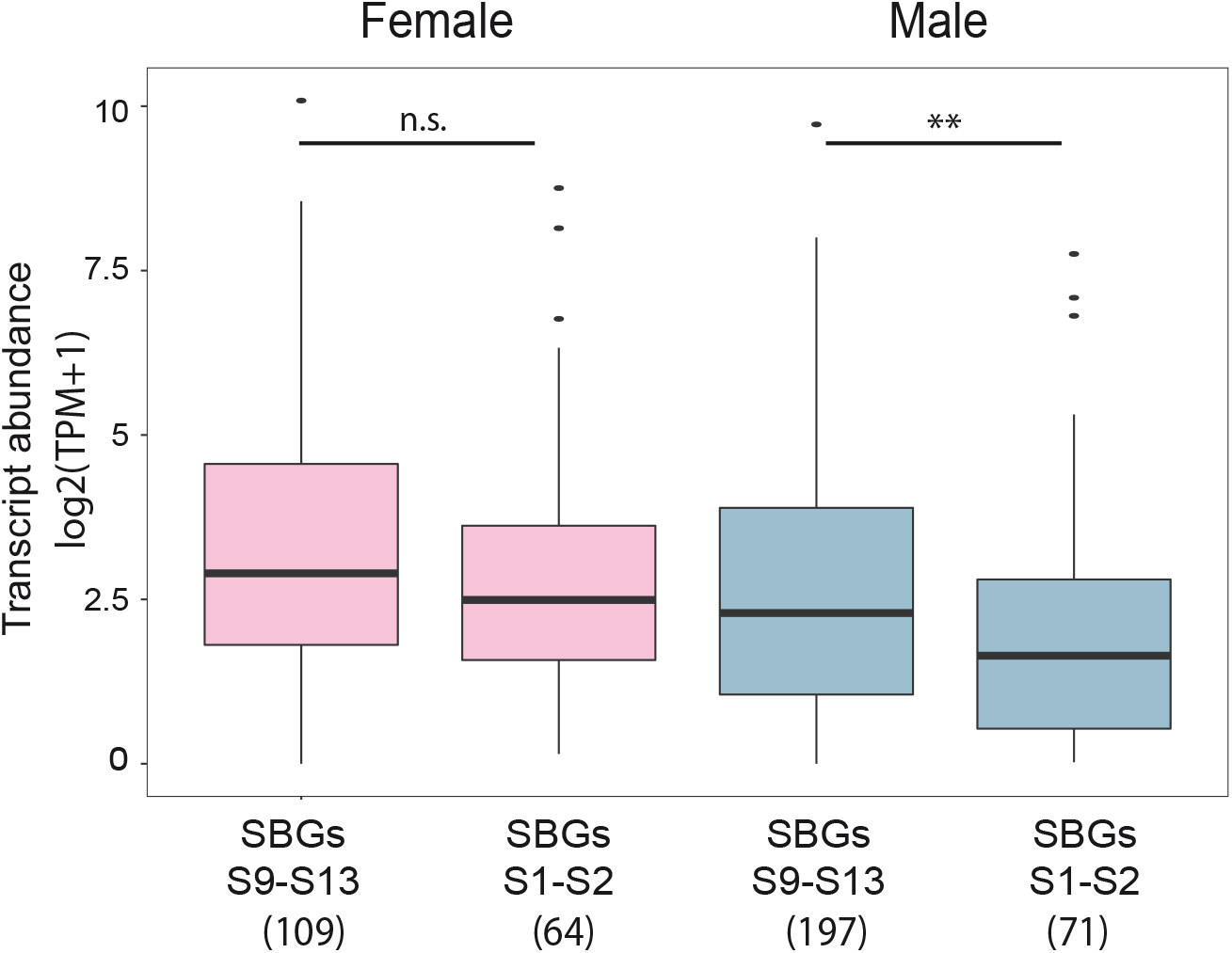
Abundances of the transcripts of sex-biased genes (SBG) marked with different chromatin states in females and males. Abundances of transcripts of SBGs in chromatin states S9-S13 or S1-S2 in females (pink) and males (blue). Values in brackets indicate the number of genes analysed. Asterisks above the plots indicate significant differences (pair-wise Wilcoxon test, **p-value<0.01).

**Figure S3.**
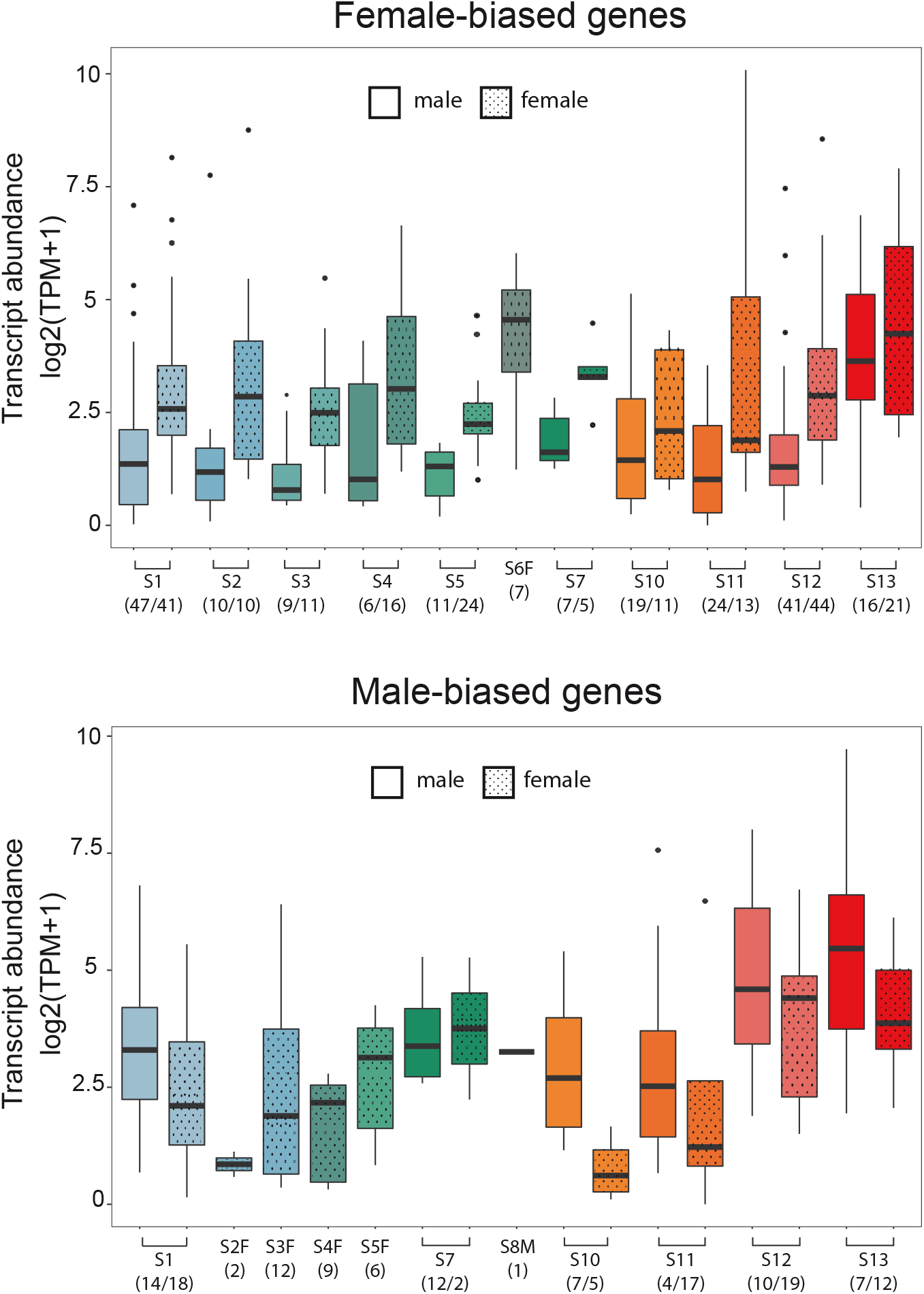
Abundances of transcripts of SBGs associated with each of the different chromatin states in males and females. The colour code is the same as that used in Figure 1A. The total number of SBGs associated with each state are indicated in brackets.

**Figure S4.**
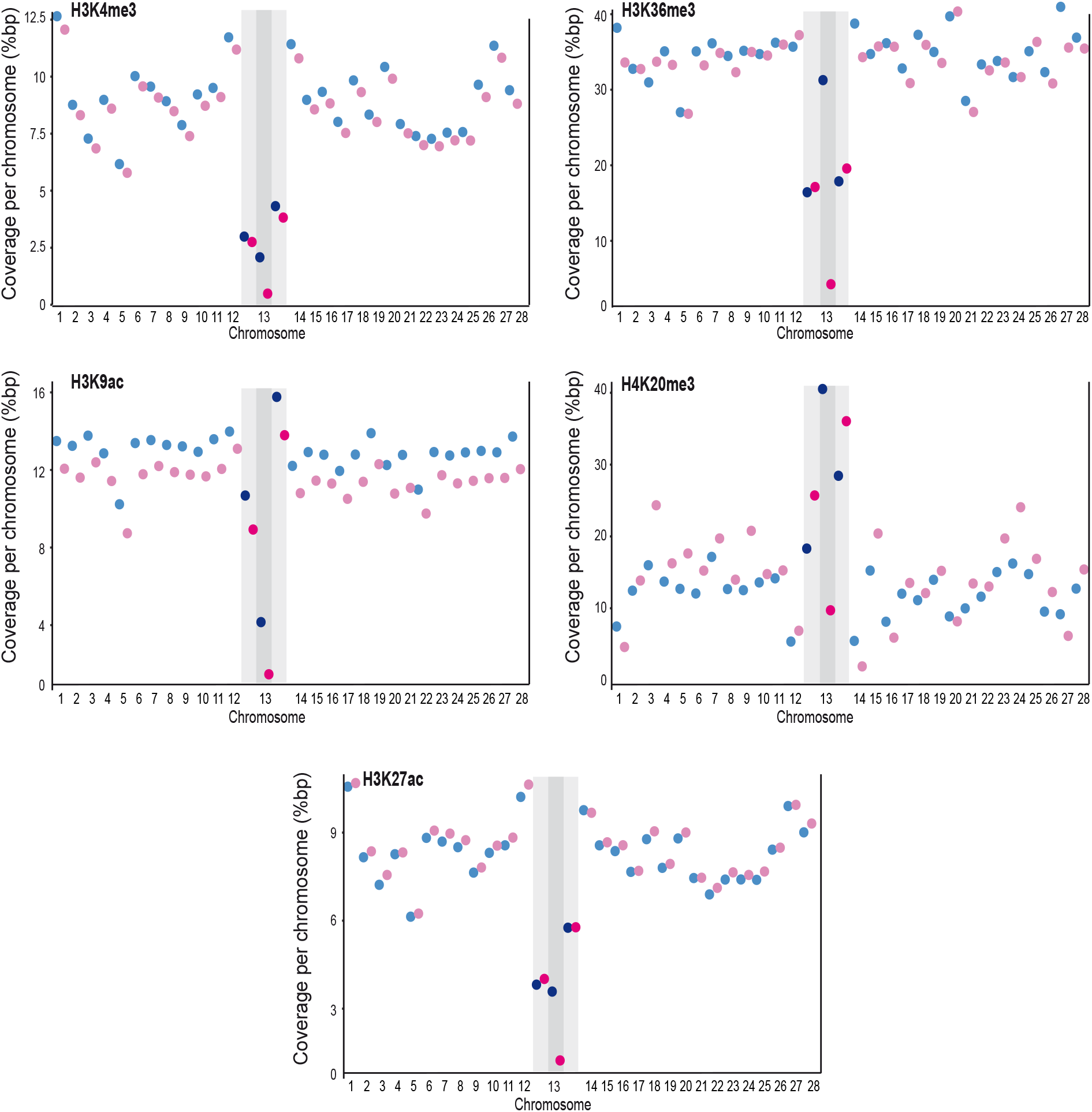
Percentage of coverage for specific histone PTMs for the SDRs, PAR and autosomes in male and females. Scatter plot showing the percent of coverage (in base pairs) for each of the five histone PTMs, H3K4me3, H3K9ac, H3K27ac, H3K36me3 and H4K20me3. Light blue and light pink represent coverage in male and female, respectively. Dark blue and red dots correspond to coverage for the V and U sex chromosomes, respectively. Light shading indicates the two PARs and dark shading the non-recombining, sex specific region (SDR) of the sex chromosome (chromosome 13).

**Figure S5.**
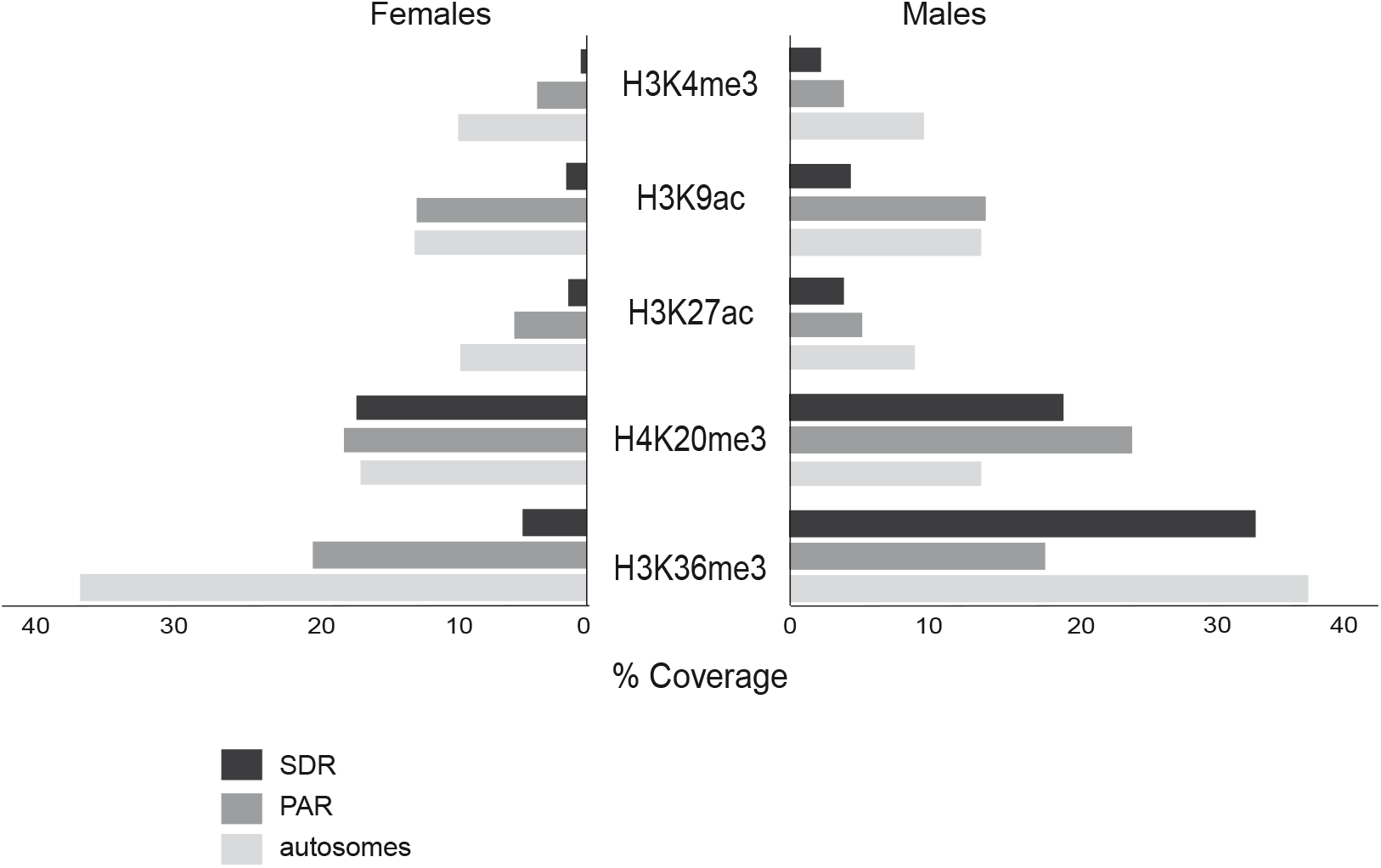
Coverage (represented as percentage of base pairs) in three different genomic regions (PAR, SDR and autosomes) marked with different histone PTMs in females (left) and males (right).

**Figure S6.**
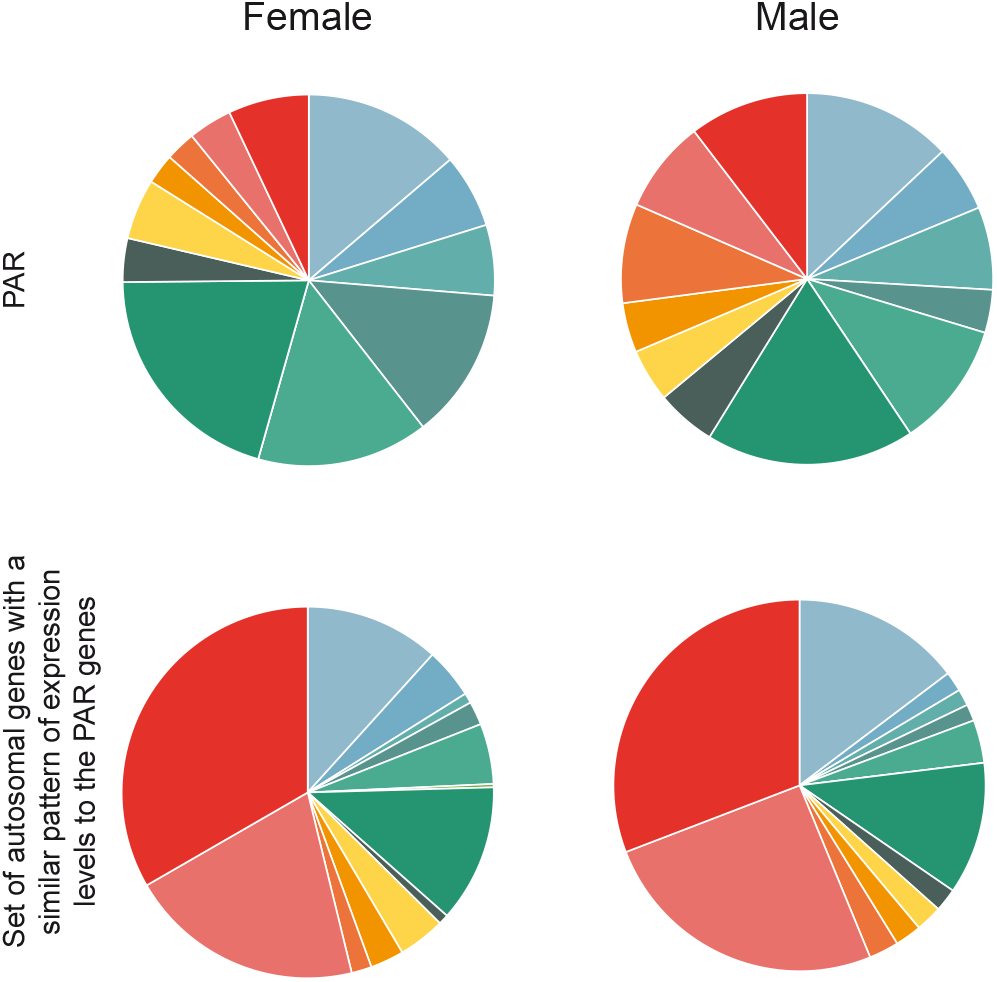
Proportions of chromatin states for PAR genes compared with the proportions of chromatin states for a set of autosomal genes with a similar pattern of expression levels to the PAR genes.

**Figure S7.**
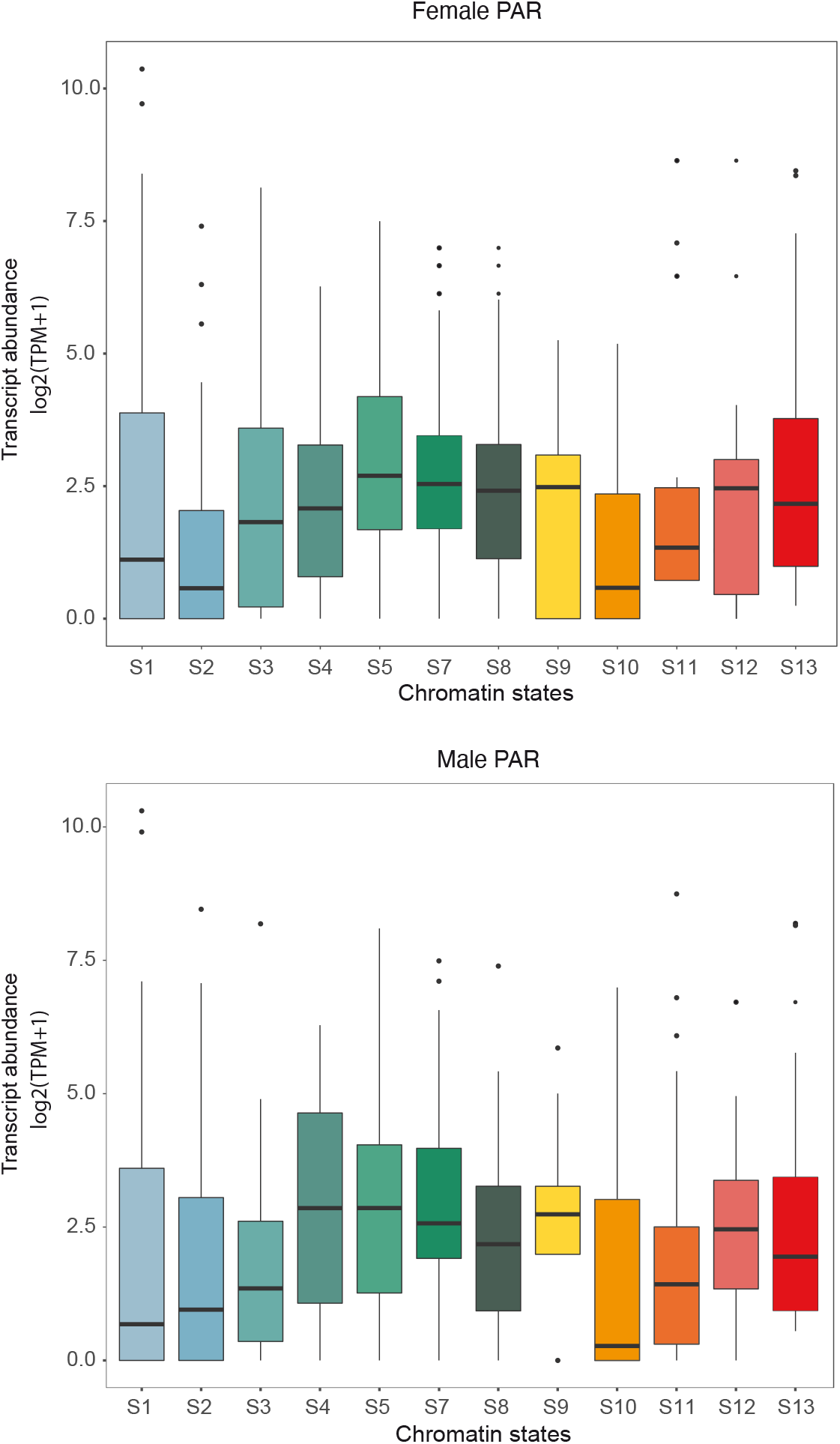
Transcript abundances, measured as log2(TPM+1), for PAR genes associated with different chromatin states in males and females. The colour code is the same as that used in Figure 1A.

**Figure S8.**
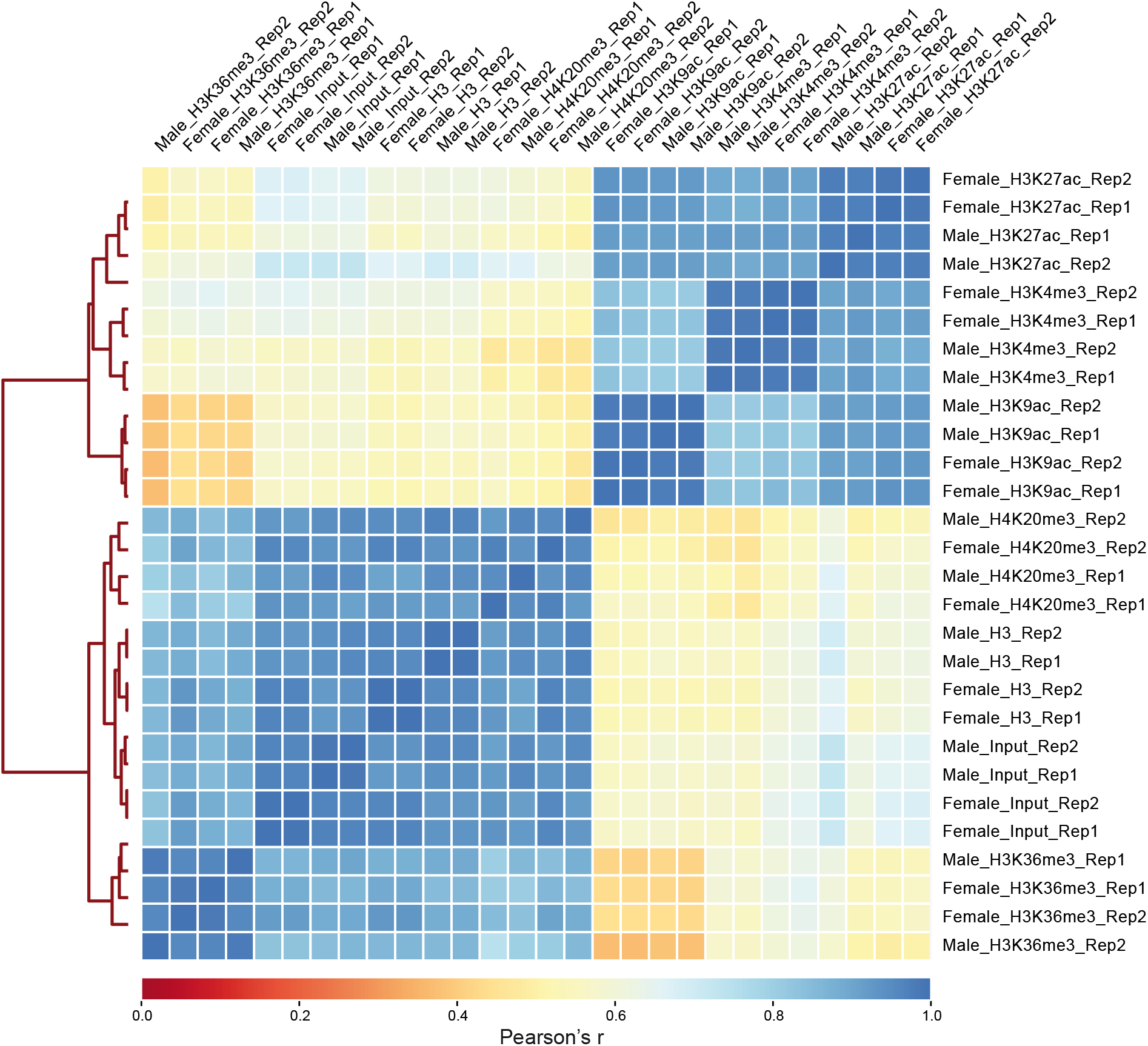
Pearson correlation scores for comparisons of the genomic distributions of ChIP-seq signal peaks for the five histone PTMs. Rep1, replicate 1; Rep2, replicate 2.

## Supplemental Tables Legends

**Table S1. *Ectocarpus* strains used, RNA-seq sequencing statistics and SRA accession numbers.**

**Table S2. Sequencing statistics for the ChIP-seq analysis and GEO reference for the dataset**. N. peaks, number of peaks; FRiP, fraction of reads in peaks.

**Table S3. Percentages of genes associated with each of the 13 chromatin states for different gene sets in males and females.** Global, all genes in the genome; Transcribed genes, genes with TPM >5th percentile; Silent genes, genes with TPM <5th percentile; Housekeeping and Narrowly-expressed, genes with tau <0.75 and tau >0.75, respectively; Unbiased, no sex-biased expression. For the chromatin states, refer to Figure 1A.

**Table S4. Percentages of narrowly expressed genes (NEG) and housekeeping (broadly expressed) genes marked with different histone PTMs in males and females.**

**Table S5. Chromatin states (S1-S13) and transcript abundances (measured as TPM) for all *Ectocarpus* genes in males and females.** FBG, female-biased gene; MBG, male-biased gene. For the chromatin states, refer to Figure 1A. Genes that did not pass the manual inspection (see methods) and were excluded from the analysis of chromatin state transitions are marked in grey.

**Table S6. Pairwise Wilcoxon tests for statistical differences between the expression levels of genes associated with specific chromatin states (Figure 2A).** S, Chromatin states (S1 to S13); F, females; M, males. The values indicate pairwise Wilcoxon test p-values corrected for multiple comparisons. For the chromatin states, refer to Figure 1A.

**Table S7. GO term enrichment for genes associated with each of the chromatin states in males and females.** All significantly enriched Biological Process GO terms identified using Blast2GO are presented.

**Table S8. Number of sex-biased genes in each of the chromatin states S1-S13 in males and females.** FBG, female-biased gene, MBG, male-biased gene

**Table S9. Transitions between chromatin states observed for male-biased and female-biased genes in males compared with females.** For chromatin states, refer to Figure 1A.

**Table S10. Coverage of the five histone PTMs across male and female genomes.** The sex chromosome (chromosome 13) is divided into PAR1 (pseudo-autosomal region 1), SDR (sex-determining region) and PAR2 (pseudo-autosomal region 2).

**Table S11. Permutation tests performed to determine whether the relative proportions of the different chromatin states were statistically different in different regions of the genome.** We randomized the genomic location of autosomal genes 100,000 times and tested the difference between the observed proportions for the SDR, the PAR or the entire sex chromosome and the permuted gene sets using chi-square statistics. Tests were performed independently for each chromatin state. Significant p-values (<0.05) are highlighted in bold.

**Table S12. Chromatin states of PAR genes in males and females.**

**Table S13. Chromatin states and transcript abundances (log2TPM+1) for SDR genes** (see also Figure 4F).

**Table S14. The presence of transposon sequences in the majority (80%) of PAR genes does not explain the distinct chromatin landscape of the PAR.** Correlation between the presence of transposable elements within introns and the presence of H4K20me3 in PAR genes and autosomal genes (left table). Permutation tests comparing the proportion of each chromatin state in the PAR with the proportion of that state in 100,000 samples of 430 autosomal genes with transposon sequences in 80% of the genes. For most chromatin states, the proportion on the PAR was significantly different from those of the autosomal gene samples indicating that transposon content does not explain the unusual pattern of chromatin states observed for the PAR. Significant p-values (<0.05) are highlighted in bold (right table).

**Table S15. Comparison of chromatin states of the PAR genes with those of a set of autosomal genes with a similar pattern of gene expression levels.** To establish the autosomal gene set, for each PAR gene, the full set of autosomal genes was searched for the gene that had the most similar level of expression. When the TPM of the PAR gene was zero, an autosomal gene with a TPM of zero was selected at random. Figure S6 presents the proportions of chromatin states associated with the two gene sets.

**Table S16. Linear models to test whether there was a significant correlation between expression level (log2(TPM + 1)) and chromatin state (upper table) or to test whether location of a gene on the PAR or on an autosome significantly influenced the expression level associated with each chromatin state (bottom table).** Significant interaction terms, in bold, represent a significantly different effect of the chromatin state on gene expression level in the PAR region compared to autosomal genes. None of the interaction terms between SDR and chromatin state showed a significant effect on gene expression so they are not reported in the table (likelihood ratio test; p-value=0.460 and p-value 0.304 for female and male SDR respectively).

## Acknowledgements

We thank Carl Herrmann and Swann Floc’hlay for advice about ChIP-seq analysis, Thomas Broquet for discussions on the statistical analysis and Maxim Bruto for help with the Circos visualisation. We thank the Institut Français de Bioinformatique and the Roscoff Analysis and Bioinformatics for Marine Science platform ABiMS (http://abims.sb-roscoff.fr) for providing computing and data storage resources. This work was supported by the CNRS, Sorbonne Université, an ERC starting grant to S.M.C. (638240) and the Agence Nationale de la Recherche project Epicycle (ANR-19-CE20-0028-01).

## References

Ahmed S, Cock JM, Pessia E, Luthringer R, Cormier A, Robuchon M, Sterck L, Peters AF, Dittami SM, Corre E, Valero M, Aury J-M, Roze D, Van de Peer Y, Bothwell J, Marais GAB, Coelho SM. 2014. A haploid system of sex determination in the brown alga Ectocarpus sp. Curr Biol 24:1945–1957. doi:10.1016/j.cub.2014.07.042

Alexa A, Rahnenfuhrer J. 2020. Enrichment Analysis for Gene Ontology.

Allis CD, Jenuwein T. 2016. The molecular hallmarks of epigenetic control. Nature Reviews Genetics 17:487.

Andrews S. 2016. FastQC A Quality Control tool for High Throughput Sequence Data. http://www.bioinformatics.babraham.ac.uk/projects/fastqc/.

Avia K, Lipinska AP, Mignerot L, Montecinos AE, Jamy M, Ahmed S, Valero M, Peters AF, Cock JM, Roze D, Coelho SM. 2018. Genetic diversity in the UV sex chromosomes of the brown alga *Ectocarpus*. Genes (Basel) 9. doi:10.3390/genes9060286

Bachtrog D. 2013. Y chromosome evolution: emerging insights into processes of Y chromosome degeneration. Nature reviews Genetics 14:113–124. doi:10.1038/nrg3366

Bachtrog D. 2006. A dynamic view of sex chromosome evolution. Curr Opin Genet Dev 16:578–585. doi:10.1016/j.gde.2006.10.007

Bachtrog D, Mank JE, Peichel CL, Kirkpatrick M, Otto SP, Ashman T-L, Hahn MW, Kitano J, Mayrose I, Ming R, Perrin N, Ross L, Valenzuela N, Vamosi JC. 2014. Sex determination: why so many ways of doing it? PLoS Biol 12:e1001899. doi:10.1371/journal.pbio.1001899

Baker BS, Gorman M, Marin I. 1994. Dosage compensation in Drosophila. Annu Rev Genet 28:491–521. doi:10.1146/annurev.ge.28.120194.002423

Baroux C, Raissig MT, Grossniklaus U. 2011. Epigenetic regulation and reprogramming during gamete formation in plants. Current Opinion in Genetics & Development 21:124–133. doi:10.1016/j.gde.2011.01.017

Barski A, Cuddapah S, Cui K, Roh T-Y, Schones DE, Wang Z, Wei G, Chepelev I, Zhao K. 2007. High-resolution profiling of histone methylations in the human genome. Cell 129:823–837. doi:10.1016/j.cell.2007.05.009

Bell O, Conrad T, Kind J, Wirbelauer C, Akhtar A, Schubeler D. 2008. Transcription-coupled methylation of histone H3 at lysine 36 regulates dosage compensation by enhancing recruitment of the MSL complex in *Drosophila melanogaster*. Mol Cell Biol 28:3401–3409. doi:10.1128/MCB.00006-08

Bisoni L, Batlle-Morera L, Bird AP, Suzuki M, McQueen HA. 2005. Female-specific hyperacetylation of histone H4 in the chicken Z chromosome. Chromosome Res 13:205–214. doi:10.1007/s10577-005-1505-4

Bourdareau S, Tirichine L, Lombard B, Loew D, Scornet D, Coelho SM, Cock JM. 2020. Histone modifications during the life cycle of the brown alga *Ectocarpus*. bioRxiv 2020.03.09.980763. doi:10.1101/2020.03.09.980763

Brockdorff N, Turner BM. 2015. Dosage compensation in mammals. Cold Spring Harb Perspect Biol 7:a019406. doi:10.1101/cshperspect.a019406

Brown EJ, Bachtrog D. 2014. The chromatin landscape of Drosophila: comparisons between species, sexes, and chromosomes. Genome Res 24:1125–1137. doi:10.1101/gr.172155.114

Brusslan JA, Bonora G, Rus-Canterbury AM, Tariq F, Jaroszewicz A, Pellegrini M. 2015. A Genome-Wide Chronological Study of Gene Expression and Two Histone Modifications, H3K4me3 and H3K9ac, during Developmental Leaf Senescence. Plant Physiol 168:1246–1261. doi:10.1104/pp.114.252999

Bull JJ. 1978. Sex Chromosomes in Haploid Dioecy: A Unique Contrast to Muller’s Theory for Diploid Dioecy. The American Naturalist 112:245–250. doi:10.1086/283267

Charlesworth B, Charlesworth D. 2000. The degeneration of Y chromosomes. Philos Trans R Soc Lond B Biol Sci 355:1563–1572. doi:10.1098/rstb.2000.0717

Charlesworth D. 2017. Evolution of recombination rates between sex chromosomes. Philos Trans R Soc Lond B Biol Sci 372. doi:10.1098/rstb.2016.0456

Cock JM, Sterck L, Rouzé P, Scornet D, Allen AE, Amoutzias G, Anthouard V, Artiguenave F, Aury J-M, Badger JH, Beszteri B, Billiau K, Bonnet E, Bothwell JH, Bowler C, Boyen C, Brownlee C, Carrano CJ, Charrier B, Cho GY, Coelho SM, Collén J, Corre E, Da Silva C, Delage L, Delaroque N, Dittami SM, Doulbeau S, Elias M, Farnham G, Gachon CMM, Gschloessl B, Heesch S, Jabbari K, Jubin C, Kawai H, Kimura K, Kloareg B, Küpper FC, Lang D, Le Bail A, Leblanc C, Lerouge P, Lohr M, Lopez PJ, Martens C, Maumus F, Michel G, Miranda-Saavedra D, Morales J, Moreau H, Motomura T, Nagasato C, Napoli CA, Nelson DR, Nyvall-Collén P, Peters AF, Pommier C, Potin P, Poulain J, Quesneville H, Read B, Rensing SA, Ritter A, Rousvoal S, Samanta M, Samson G, Schroeder DC, Ségurens B, Strittmatter M, Tonon T, Tregear JW, Valentin K, von Dassow P, Yamagishi T, Van de Peer Y, Wincker P. 2010. The *Ectocarpus* genome and the independent evolution of multicellularity in brown algae. Nature 465:617–621. doi:10.1038/nature09016

Coelho SM, Gueno J, Lipinska AP, Cock JM, Umen JG. 2018. UV chromosomes and haploid sexual systems. Trends Plant Sci 23:794–807. doi:10.1016/j.tplants.2018.06.005

Coelho SM, Mignerot L, Cock JM. 2019. Origin and evolution of sex-determination systems in the brown algae. New Phytologist **10.1111/nph.15694**. doi:10.1111/nph.15694

Coelho SM, Scornet D, Rousvoal S, Peters NT, Dartevelle L, Peters AF, Cock JM. 2012. How to cultivate *Ectocarpus*. Cold Spring Harb Protoc 2012:258–261. doi:10.1101/pdb.prot067934

Cormier A, Avia K, Sterck L, Derrien T, Wucher V, Andres G, Monsoor M, Godfroy O, Lipinska A, Perrineau M-M, Van De Peer Y, Hitte C, Corre E, Coelho SM, Cock JM. 2017a. Re-annotation, improved large-scale assembly and establishment of a catalogue of noncoding loci for the genome of the model brown alga *Ectocarpus*. New Phytol 214:219–232. doi:10.1111/nph.14321

Cormier A, Avia K, Sterck L, Derrien T, Wucher V, Andres G, Monsoor M, Godfroy O, Lipinska A, Perrineau M-M, Van De Peer Y, Hitte C, Corre E, Coelho SM, Cock JM. 2017b. Re-annotation, improved large-scale assembly and establishment of a catalogue of noncoding loci for the genome of the model brown alga *Ectocarpus*. New Phytol 214:219–232. doi:10.1111/nph.14321

Creyghton MP, Cheng AW, Welstead GG, Kooistra T, Carey BW, Steine EJ, Hanna J, Lodato MA, Frampton GM, Sharp PA, Boyer LA, Young RA, Jaenisch R. 2010. Histone H3K27ac separates active from poised enhancers and predicts developmental state. Proceedings of the National Academy of Sciences 107:21931–21936. doi:10.1073/pnas.1016071107

de la Paz Sanchez M, Gutierrez C. 2009. *Arabidopsis* ORC1 is a PHD-containing H3K4me3 effector that regulates transcription. Proc Natl Acad Sci USA 106:2065. doi:10.1073/pnas.0811093106

Diaz A, Park K, Lim DA, Song JS. 2012. Normalization, bias correction, and peak calling for ChIP-seq. Stat Appl Genet Mol Biol 11:Article 9. doi:10.1515/1544-6115.1750

Elango N, Hunt BG, Goodisman MAD, Yi SV. 2009. DNA methylation is widespread and associated with differential gene expression in castes of the honeybee, Apis mellifera. Proc Natl Acad Sci U S A 106:11206–11211. doi:10.1073/pnas.0900301106

Filion GJ, van Bemmel JG, Braunschweig U, Talhout W, Kind J, Ward LD, Brugman W, de Castro IJ, Kerkhoven RM, Bussemaker HJ, van Steensel B. 2010. Systematic protein location mapping reveals five principal chromatin types in Drosophila cells. Cell 143:212–224. doi:10.1016/j.cell.2010.09.009

Fischer A, Hofmann I, Naumann K, Reuter G. 2006. Heterochromatin proteins and the control of heterochromatic gene silencing in Arabidopsis. J Plant Physiol 163:358–368. doi:10.1016/j.jplph.2005.10.015

Gelbart ME, Kuroda MI. 2009. Drosophila dosage compensation: a complex voyage to the X chromosome. Development 136:1399–1410. doi:10.1242/dev.029645

Girton JR, Johansen KM. 2008. Chromatin structure and the regulation of gene expression: the lessons of PEV in Drosophila. Adv Genet 61:1–43. doi:10.1016/S0065-2660(07)00001-6

Grath S, Parsch J. 2016. Sex-Biased Gene Expression. Annu Rev Genet 50:29–44. doi:10.1146/annurev-genet-120215-035429

Hansen P, Hecht J, Ibn-Salem J, Menkuec BS, Roskosch S, Truss M, Robinson PN. 2016. Q-nexus: a comprehensive and efficient analysis pipeline designed for ChIP-nexus. BMC Genomics 17:873. doi:10.1186/s12864-016-3164-6

He G, Zhu X, Elling AA, Chen L, Wang X, Guo L, Liang M, He H, Zhang H, Chen F, Qi Y, Chen R, Deng X-W. 2010. Global epigenetic and transcriptional trends among two rice subspecies and their reciprocal hybrids. Plant Cell 22:17–33. doi:10.1105/tpc.109.072041

Heintzman ND, Stuart RK, Hon G, Fu Y, Ching CW, Hawkins RD, Barrera LO, Van Calcar S, Qu C, Ching KA, Wang W, Weng Z, Green RD, Crawford GE, Ren B. 2007. Distinct and predictive chromatin signatures of transcriptional promoters and enhancers in the human genome. Nat Genet 39:311–318. doi:10.1038/ng1966

Howe FS, Fischl H, Murray SC, Mellor J. 2017. Is H3K4me3 instructive for transcription activation? Bioessays 39:1–12. doi:10.1002/bies.201600095

Huang C, Zhu B. 2018. Roles of H3K36-specific histone methyltransferases in transcription: antagonizing silencing and safeguarding transcription fidelity. Biophys Rep 4:170–177. doi:10.1007/s41048-018-0063-1

Jones PA. 2012. Functions of DNA methylation: islands, start sites, gene bodies and beyond. Nat Rev Genet 13:484–492. doi:10.1038/nrg3230

Kim D, Pertea G, Trapnell C, Pimentel H, Kelley R, Salzberg SL. 2013. TopHat2: accurate alignment of transcriptomes in the presence of insertions, deletions and gene fusions. Genome Biol 14:R36. doi:10.1186/gb-2013-14-4-r36

Kouzarides T. 2007. Chromatin modifications and their function. Cell 128:693–705. doi:10.1016/j.cell.2007.02.005

Krzywinski M, Schein J, Birol I, Connors J, Gascoyne R, Horsman D, Jones SJ, Marra MA. 2009. Circos: an information aesthetic for comparative genomics. Genome Res 19:1639–1645. doi:10.1101/gr.092759.109

Landt SG, Marinov GK, Kundaje A, Kheradpour P, Pauli F, Batzoglou S, Bernstein BE, Bickel P, Brown JB, Cayting P, Chen Y, DeSalvo G, Epstein C, Fisher-Aylor KI, Euskirchen G, Gerstein M, Gertz J, Hartemink AJ, Hoffman MM, Iyer VR, Jung YL, Karmakar S, Kellis M, Kharchenko PV, Li Q, Liu T, Liu XS, Ma L, Milosavljevic A, Myers RM, Park PJ, Pazin MJ, Perry MD, Raha D, Reddy TE, Rozowsky J, Shoresh N, Sidow A, Slattery M, Stamatoyannopoulos JA, Tolstorukov MY, White KP, Xi S, Farnham PJ, Lieb JD, Wold BJ, Snyder M. 2012. ChIP-seq guidelines and practices of the ENCODE and modENCODE consortia. Genome Research 22:1813–1831. doi:10.1101/gr.136184.111

Langmead B, Trapnell C, Pop M, Salzberg SL. 2009. Ultrafast and memory-efficient alignment of short DNA sequences to the human genome. Genome Biol 10:R25. doi:10.1186/gb-2009-10-3-r25

Lemos B, Branco AT, Hartl DL. 2010. Epigenetic effects of polymorphic Y chromosomes modulate chromatin components, immune response, and sexual conflict. Proc Natl Acad Sci U S A 107:15826–15831. doi:10.1073/pnas.1010383107

Lerdrup M, Johansen JV, Agrawal-Singh S, Hansen K. 2016. An interactive environment for agile analysis and visualization of ChIP-sequencing data. Nat Struct Mol Biol 23:349–357. doi:10.1038/nsmb.3180

Lesch BJ, Page DC. 2013. Sex-specific chromatin states in mammalian fetal germ cells. Epigenetics Chromatin 6:P45–P45. doi:10.1186/1756-8935-6-S1-P45

Li H, Handsaker B, Wysoker A, Fennell T, Ruan J, Homer N, Marth G, Abecasis G, Durbin R, 1000 Genome Project Data Processing Subgroup. 2009. The Sequence Alignment/Map format and SAMtools. Bioinformatics 25:2078–2079. doi:10.1093/bioinformatics/btp352

Liao Y, Smyth GK, Shi W. 2014. featureCounts: an efficient general purpose program for assigning sequence reads to genomic features. Bioinformatics 30:923–930. doi:10.1093/bioinformatics/btt656

Lindeman LC, Winata CL, Aanes H, Mathavan S, Alestrom P, Collas P. 2010. Chromatin states of developmentally-regulated genes revealed by DNA and histone methylation patterns in zebrafish embryos. Int J Dev Biol 54:803–813. doi:10.1387/ijdb.103081ll

Lipinska AP, D’hondt S, Van Damme EJ, De Clerck O. 2013. Uncovering the genetic basis for early isogamete differentiation: a case study of *Ectocarpus siliculosus*. BMC Genomics 14:909–909. doi:10.1186/1471-2164-14-909

Lipinska AP, Serrano-Serrano ML, Cormier A, Peters AF, Kogame K, Cock JM, Coelho SM. 2019. Rapid turnover of life-cycle-related genes in the brown algae. Genome Biol 20:35. doi:10.1186/s13059-019-1630-6

Lipinska AP, Toda NRT, Heesch S, Peters AF, Cock JM, Coelho SM. 2017. Multiple gene movements into and out of haploid sex chromosomes. Genome Biol 18:104. doi:10.1186/s13059-017-1201-7

Lipinska, Cormier A, Luthringer R, Peters AF, Corre E, Gachon CMM, Cock JM, Coelho SM. 2015. Sexual dimorphism and the evolution of sex-biased gene expression in the brown alga *Ectocarpus*. Mol Biol Evol 32:1581–1597. doi:10.1093/molbev/msv049

Love MI, Huber W, Anders S. 2014. Moderated estimation of fold change and dispersion for RNA-seq data with DESeq2. Genome Biol 15:550. doi:10.1186/s13059-014-0550-8

Lucchesi JC, Kelly WG, Panning B. 2005. Chromatin remodeling in dosage compensation. Annu Rev Genet 39:615–651. doi:10.1146/annurev.genet.39.073003.094210

Luthringer R, Lipinska AP, Roze D, Cormier A, Macaisne N, Peters AF, Cock JM, Coelho SM. 2015. The pseudoautosomal regions of the U/V sex chromosomes of the brown alga *Ectocarpus* exhibit unusual features. Mol Biol Evol 32:2973–2985. doi:10.1093/molbev/msv173

Margueron R, Reinberg D. 2010. Chromatin structure and the inheritance of epigenetic information. Nat Rev Genet 11:285–296. doi:10.1038/nrg2752

Martin M. 2011. Cutadapt removes adapter sequences from high-throughput sequencing reads. EMBnet 10–12. doi:https://doi.org/10.14806/ej.17.1.200

Mignerot L, Coelho SM. 2016. The origin and evolution of the sexes: Novel insights from a distant eukaryotic linage. C R Biol 339:252–257. doi:10.1016/j.crvi.2016.04.012

Nelson DM, Jaber-Hijazi F, Cole JJ, Robertson NA, Pawlikowski JS, Norris KT, Criscione SW, Pchelintsev NA, Piscitello D, Stong N, Rai TS, McBryan T, Otte GL, Nixon C, Clark W, Riethman H, Wu H, Schotta G, Garcia BA, Neretti N, Baird DM, Berger SL, Adams PD. 2016. Mapping H4K20me3 onto the chromatin landscape of senescent cells indicates a function in control of cell senescence and tumor suppression through preservation of genetic and epigenetic stability. Genome Biology 17:158. doi:10.1186/s13059-016-1017-x

Nugent BM, Wright CL, Shetty AC, Hodes GE, Lenz KM, Mahurkar A, Russo SJ, Devine SE, McCarthy MM. 2015. Brain feminization requires active repression of masculinization via DNA methylation. Nat Neurosci 18:690–697. doi:10.1038/nn.3988

Picard MAL, Cosseau C, Ferre S, Quack T, Grevelding CG, Coute Y, Vicoso B. 2018. Evolution of gene dosage on the Z-chromosome of schistosome parasites. Elife 7. doi:10.7554/eLife.35684

Picard MAL, Vicoso B, Roquis D, Bulla I, Augusto RC, Arancibia N, Grunau C, Boissier J, Cosseau C. 2019. Dosage Compensation throughout the Schistosoma mansoni Lifecycle: Specific Chromatin Landscape of the Z Chromosome. Genome Biol Evol 11:1909–1922. doi:10.1093/gbe/evz133

Quinlan AR, Hall IM. 2010. BEDTools: a flexible suite of utilities for comparing genomic features. Bioinformatics 26:841–842. doi:10.1093/bioinformatics/btq033

Ramirez F, Dundar F, Diehl S, Gruning BA, Manke T. 2014. deepTools: a flexible platform for exploring deep-sequencing data. Nucleic Acids Res 42:W187–191. doi:10.1093/nar/gku365

Roudier F, Ahmed I, Bérard C, Sarazin A, Mary-Huard T, Cortijo S, Bouyer D, Caillieux E, Duvernois-Berthet E, Al-Shikhley L, Giraut L, Després B, Drevensek S, Barneche F, Dèrozier S, Brunaud V, Aubourg S, Schnittger A, Bowler C, Martin-Magniette M-L, Robin S, Caboche M, Colot V. 2011. Integrative epigenomic mapping defines four main chromatin states in Arabidopsis. The EMBO Journal 30:1928–1938. doi:10.1038/emboj.2011.103

Schmid MW, Giraldo-Fonseca A, Rovekamp M, Smetanin D, Bowman JL, Grossniklaus U. 2018. Extensive epigenetic reprogramming during the life cycle of Marchantia polymorpha. Genome Biol 19:9. doi:10.1186/s13059-017-1383-z

Schotta G, Lachner M, Sarma K, Ebert A, Sengupta R, Reuter G, Reinberg D, Jenuwein T. 2004. A silencing pathway to induce H3-K9 and H4-K20 trimethylation at constitutive heterochromatin. Genes Dev 18:1251–1262. doi:10.1101/gad.300704

She W, Baroux C. 2015. Chromatin dynamics in pollen mother cells underpin a common scenario at the somatic-to-reproductive fate transition of both the male and female lineages in Arabidopsis. Front Plant Sci 6:294. doi:10.3389/fpls.2015.00294

Shilatifard A. 2006. Chromatin modifications by methylation and ubiquitination: implications in the regulation of gene expression. Annu Rev Biochem 75:243–269. doi:10.1146/annurev.biochem.75.103004.142422

Srivastava S, Mishra RK, Dhawan J. 2010. Regulation of cellular chromatin state: insights from quiescence and differentiation. Organogenesis 6:37–47. doi:10.4161/org.6.1.11337

Straub T, Becker PB. 2007. Dosage compensation: the beginning and end of generalization. Nat Rev Genet 8:47–57. doi:10.1038/nrg2013

Umen J, Coelho S. 2019. Algal Sex Determination and the Evolution of Anisogamy. Annu Rev Microbiol. doi:10.1146/annurev-micro-020518-120011

Vicoso B, Charlesworth B. 2009. Effective population size and the faster-X effect: an extended model. Evolution 63:2413–2426. doi:10.1111/j.1558-5646.2009.00719.x

Xu S, Grullon S, Ge K, Peng W. 2014. Spatial clustering for identification of ChIP-enriched regions (SICER) to map regions of histone methylation patterns in embryonic stem cells. Methods Mol Biol 1150:97–111. doi:10.1007/978-1-4939-0512-6_5

Yanai I, Benjamin H, Shmoish M, Chalifa-Caspi V, Shklar M, Ophir R, Bar-Even A, Horn-Saban S, Safran M, Domany E, Lancet D, Shmueli O. 2005. Genome-wide midrange transcription profiles reveal expression level relationships in human tissue specification. Bioinformatics 21:650–659. doi:10.1093/bioinformatics/bti042

Yasuhara JC, Wakimoto BT. 2008. Molecular landscape of modified histones in Drosophila heterochromatic genes and euchromatin-heterochromatin transition zones. PLoS Genet 4:e16. doi:10.1371/journal.pgen.0040016

Zang C, Schones DE, Zeng C, Cui K, Zhao K, Peng W. 2009. A clustering approach for identification of enriched domains from histone modification ChIP-Seq data. Bioinformatics 25:1952–1958. doi:10.1093/bioinformatics/btp340

Zhang Y, Liu T, Meyer CA, Eeckhoute J, Johnson DS, Bernstein BE, Nusbaum C, Myers RM, Brown M, Li W, Liu XS. 2008. Model-based analysis of ChIP-Seq (MACS). Genome Biol 9:R137. doi:10.1186/gb-2008-9-9-r137

